# Structure and dynamics of *Burkholderia pseudomallei* OXA-57, a distinctive low efficiency class D β-lactamase with carbapenem-hydrolyzing activity

**DOI:** 10.1101/2024.12.23.630153

**Authors:** Éilís C. Bragginton, Charlotte K. Colenso, Karina Calvopiña, Philip Hinchliffe, John M. Shaw, Catherine. L. Tooke, Rathanin Seng, Narisara Chantratita, Adrian J. Mulholland, Christopher J. Schofield, James Spencer

## Abstract

The Gram-negative bacterium *Burkholderia pseudomallei* causes the severe disease melioidosis. β-Lactams, including carbapenems, are the primary treatment, but are susceptible to chromosomal β-lactamases, including the class D enzyme OXA-57. Here we show recombinant OXA-57 is active towards penicillins and first-generation cephalosporins, slowly hydrolyzes carbapenems including imipenem and meropenem, but is inactive towards oxyimino-cephalosporins (e.g., ceftazidime). Unlike many OXA enzymes, OXA-57 is sensitive to the mechanism-based inhibitor clavulanic acid, but less so to the diazabicyclooctane avibactam and not to the cyclic boronate vaborbactam. Crystal structures of covalent OXA-57:avibactam and OXA-57:meropenem complexes reveal limited hydrogen-bonding interactions with bound ligands. In molecular dynamics simulations, bound meropenem is mobile, while the water necessary for deacylation has only limited active site access. These observations are consistent with the low level of meropenem turnover, supporting proposals that OXA β-lactamases generally possess limited carbapenemase activity, and highlight the potential importance of OXA-57 in *B. pseudomallei* β-lactam resistance.

**Graphical Abstract:** 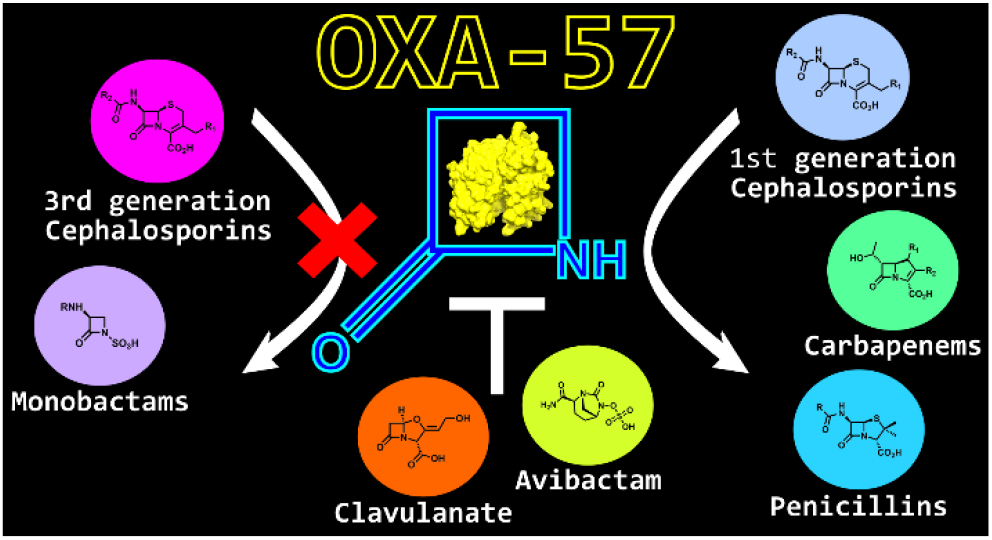

## 1. Introduction

*Burkholderia pseudomallei* is a Gram-negative intracellular pathogen that causes melioidosis, a severe disease endemic to South East Asia and Northern Australia^1^. *B. pseudomallei* infection occurs through multiple routes, including percutaneous inoculation, inhalation and ingestion^1,2^ and presents as an array of symptoms such as soft tissue lesions, fever, pneumonia and life threatening sepsis^3,4^. Melioidosis causes an estimated 89 000 deaths annually with a fatality rate of up to 50% in endemic regions^5^. The high morbidity and mortality associated with infection classes *B. pseudomallei* as a category B biological agent with potential use as a weapon in bioterrorism^6^. Survivors of acute melioidosis are susceptible to recurrent *B. pseudomallei* infection and those with pre-existing risk factors, such as diabetes mellitus, account for approximately 80% of patients^2,5^.

*B. pseudomallei* manifests intrinsic resistance to a range of antibiotics, including penicillins, 1^st^ and 2^nd^ generation cephalosporins, colistin, macrolides, aminoglycosides and rifamycins^7^. As a result, the treatment options for melioidosis are limited, with β-lactams such as carbapenems and the 3^rd^ generation cephalosporin ceftazidime being the primary therapeutic choices. Treatment involves an initial intensive phase of intravenous β-lactam antibiotics, usually meropenem or ceftazidime, over 10 - 14 days, followed by a longer course of oral antibiotics. The latter is classed as the eradication phase of treatment with therapy lasting at least 3 months using combination therapies such as trimethoprim/sulphamethoxazole or amoxicillin/clavulanate^5,7^.

To date, resistance to the clinically used β-lactam antibiotics in *B. pseudomallei* is rare. However, as with most human bacterial infections, cases of drug-resistant melioidosis are on the rise due to increased antibiotic use^8,9^. There are multiple mechanisms of resistance to β-lactam antibiotics, including efflux and decreased porin expression. However, in Gram-negative pathogens, β-lactam degradation by β-lactamases is the most common mechanism^10^. Currently, over 8,432 β-lactamases have been identified^11^; they are categorized by their mechanism of action into serine β-lactamases (SBLs) that utilize an active site serine nucleophile to catalyze β-lactam hydrolysis through formation of a covalent acyl-enzyme, and metallo β-lactamases (MBLs), where zinc ions activate a nucleophilic water molecule^12^. They can be further classified, based on amino acid sequence, into three classes of SBLs (A, C and D) and the class B MBLs^13^.

A notable property of the class-D (OXA) β-lactamases, that differentiates them from class A and C SBLs, is the requirement for a carbamylated lysine for both the acylation and deacylation steps of hydrolysis^14^ (**Figure S1**). This reversible post-translational modification is driven by reaction with atmospherically-derived CO_2_ and is facilitated by the hydrophobicity of the active site, which modifies the lysine p*K_a_* to favor carbamylation^14–16^. Some class D SBLs are also distinguished by their hydrolytic activity towards carbapenems, which more usually form long-lived inhibitory acyl-enzyme complexes on reaction with SBLs^17^. However, these carbapenem-hydrolyzing class-D β-lactamases (CHDLs) are distinguished from other SBLs with activity towards carbapenems by their employment of an alternative mechanism of carbapenem hydrolysis whereby carbapenem degradation can additionally result in the formation of β- lactone products^18^. CHDLs are important contributors to carbapenem resistance in several species of pathogenic Gram-negative bacteria, including *Acinetobacter baumannii*^19^.

Previous analysis of the *B. pseudomallei* genome identified 7 potential β-lactamases, 2 of which are described as serine-β-lactamases and 5 as putative metallo-β-lactamases^20^. To date, however, only the serine-β-lactamases PenI and OXA-57 have been characterized and shown to have β-lactamase activity^21,22^. PenI, a Class-A extended spectrum β-lactamase (ESBL), has been extensively studied ^21,23^. Previous investigations have identified PenI mutations, for example P167S^9^ and C69Y, as the cause of ceftazidime-resistant phenotypes in clinical *B. pseudomallei* isolates^8,24–26^. Similarly, the S72F PenI substitution results in resistance to the widely used β-lactamase inhibitor clavulanic acid^8,25^. However, although there is clear evidence implicating PenI as a contributing factor to ceftazidime resistance within the clinic, PenI does not catalyze carbapenem turnover^21^ and the cause of carbapenem resistance in *B. pseudomallei* clinical isolates remains elusive.

By contrast with PenI, few reported studies have investigated the possible involvement of the other validated SBL present in the *B. pseudomallei* genome, i.e. the Class-D OXA-57, which is associated with *B. pseudomallei* resistance phenotypes. A previous biochemical study of OXA-57 was limited to establishing its hydrolytic activity against penicillins and 1^st^ generation cephalosporins^22^. Accordingly, we have here undertaken a more extensive kinetic characterization of OXA-57 using a broader range of β-lactam substrates with a focus on those of current clinical relevance, as well as investigating its susceptibility to a range of chemically distinct β-lactam inhibitors, including those (avibactam, vaborbactam) recently approved for clinical use. We further present crystal structures of OXA-57 and its covalent complexes with the carbapenem meropenem and the inhibitor avibactam; and on the basis of these structural data apply extended molecular dynamics simulations to study the behavior of the meropenem derived acyl-enzyme complex.

## 2. Results

OXA-57, lacking its predicted signal peptide, was expressed in recombinant *E. coli* BL21 (DE3) using the pET28 T7 vector and was readily purified to homogeneity using immobilized metal ion affinity and size-exclusion chromatography (**Figure S2**). The purified protein eluted from the size exclusion column as a single peak at a volume consistent with the predicted molecular mass of the monomeric protein (27.1 kDa), with no evidence of a second peak that would indicate the presence of a dimer, as observed for some other class-D β-lactamases including OXA-10 and OXA-48^27–29^. While previous data indicated a propensity for OXA-57 to dimerize in solution, in that study the dimeric form represented only 10% of the total protein content^22^; hence we propose the monomer, rather than the dimer, to be the favored form of OXA-57.

### Hydrolysis of β-lactam substrates by OXA-57

Class D (OXA) β-lactamases differ widely in activity towards different β-lactams, with some (e.g. OXA-20^30^) efficiently hydrolyzing only penicillin substrates, while the activity of others extends to 3^rd^ generation cephalosporins (e.g. OXA-10^31^) and/or carbapenems (e.g. OXA-48^28^). OXA-57 was reported to turn over penicillins and 1^st^ generation cephalosporins^22^. However, the activity of OXA-57 against later-generation cephalosporins (e.g. ceftazidime) and carbapenems, and the effect of NaHCO_3_ supplementation (to ensure maximal carbamylation of the active site lysine 73) was not investigated. We therefore determined steady-state kinetic parameters for OXA-57-catalyzed hydrolysis of a wider range of β-lactam substrates in the presence of NaHCO_3_ (**Table 1**). *k_cat_*/*K*_M_ values for turnover of the aminopenicillin ampicillin and the 1^st^ generation cephalosporin cephalothin were close to those previously obtained (0.0075 and 0.0017 s^-1^ μM^-1^, respectively)^22^. However, hydrolysis was not observed for the (oxyimino)cephalosporin substrates; cefotaxime and ceftazidime, or the monobactam aztreonam. By contrast, we observed low levels of hydrolysis for the carbapenems imipenem and meropenem, but it was necessary to undertake the steady-state experiments over several hours to obtain kinetic parameters (**Figure S3a**). OXA-57 catalyzed turnover of both imipenem and meropenem was slow (*k*_cat_, 0.04 ± 0.005 and 0.01 ± 0.002 s^-1^), with *K*_M_ values (51 ± 12 and 21 ± 7 μM) ∼10-fold weaker than those typically obtained for other class D β-lactamases (**Table S2**). In comparison with other class D β-lactamases, these values are closest to those described for OXA-10, an enzyme with only weak activity towards carbapenems that is not usually considered a CHDL^32^, rather than for members of the main CHDL lineages e.g. OXA- 23 and OXA-48 (**Table S2**). Substrate inhibition of OXA-57 by both carbapenems was observed at higher concentrations (**Figure S3b**).

**Table 1.**
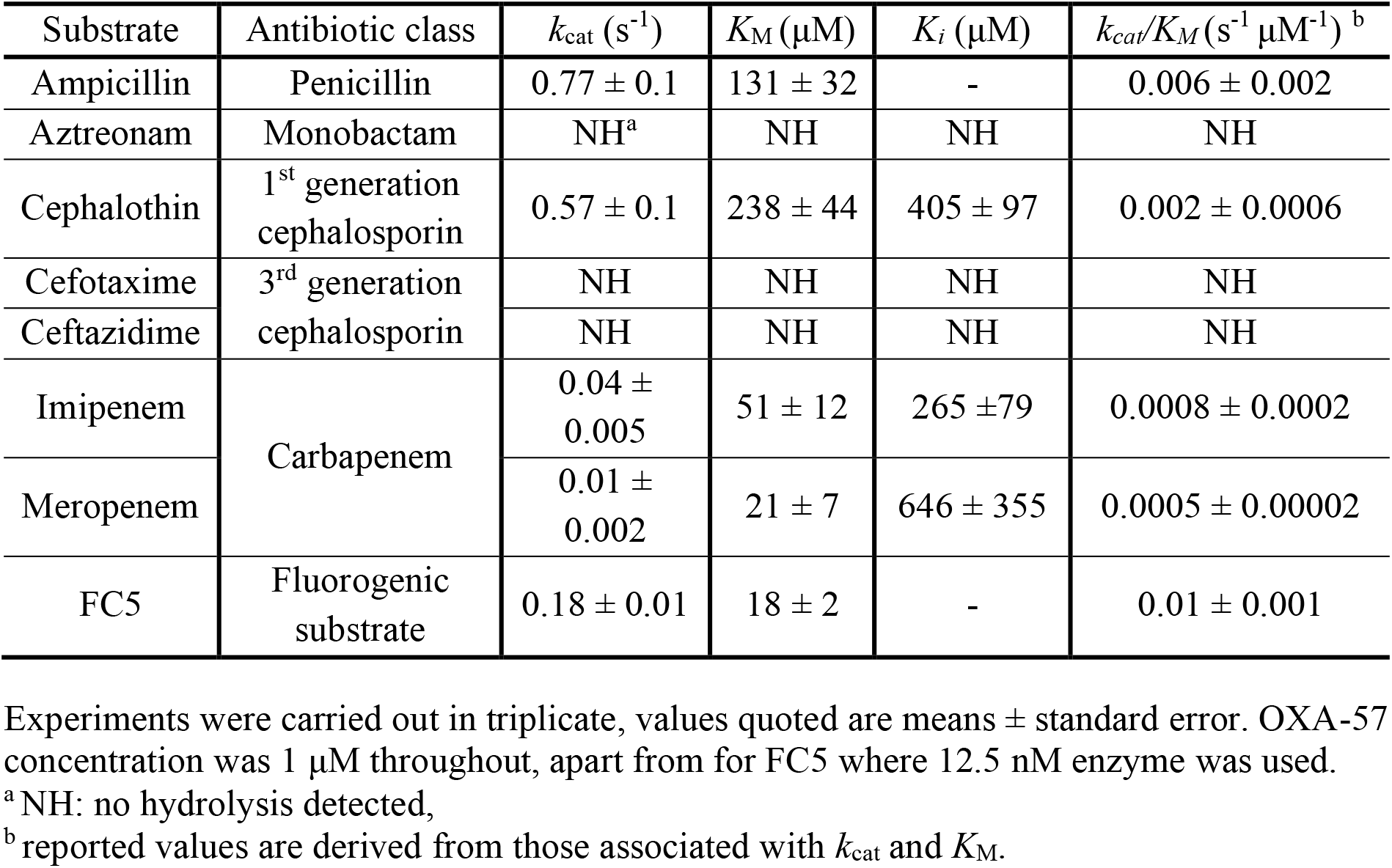
Steady-state kinetic parameters for hydrolysis of selected β-lactams by purified recombinant OXA-57.

### Inhibition of OXA-57

β-Lactamase inhibitors (BLIs) are used in combination with β-lactams to circumvent β- lactamase activity, so extending the clinically useful lifetime of β-lactams^33^. Both β-lactam compounds (e.g. clavulanate and tazobactam) that act as mechanism-based inhibitors, and non-classical inhibitors lacking a β-lactam core (e.g. the diazabicyclooctane (DBO) avibactam and the cyclic boronate vaborbactam) are now in clinical use. While clavulanate has previously been reported to inhibit OXA-57, (*K_i_* 3.4 μM)^22^, its susceptibility to the new inhibitor classes is unreported. We measured half maximal inhibitory concentrations (IC_50_) for inhibition of OXA-57 by clavulanate, avibactam and vaborbactam, measuring residual activity after pre-incubation with inhibitor for 10, 30 and 60 minutes (**Table 2**). Direct competition assays were carried out in the presence of NaHCO_3_, using as a reporter substrate the fluorogenic cephalosporin FC5^34^, which showed comparable kinetic parameters to other OXA-57 substrates (**Table 1**). Clavulanate was the most effective OXA-57 inhibitor tested, with IC_50_ values ranging from 9.4 - 30.3 μM observed, depending upon pre-incubation time, and comparable to those for other class-D β-lactamases such as OXA-10 or OXA-48 (**Table S3**). Avibactam has slightly higher IC_50_ values, from 13.2 - 58.8 μM (**Table 1**), and is a less efficient inhibitor of OXA-57 than of some other OXA enzymes, e.g. OXA-10, OXA-23 and OXA-48, for which IC_50_ values in the nM range have been reported (**Table S3**). Vaborbactam only very weakly inhibited OXA-57, consistent with its reported activity towards other class-D β- lactamases^35^ (**Table S3**), with an IC_50_ unable to be determined from measurements at up to 4 mM inhibitor.

**Table 2.** pIC_50_ and IC_50_ values for inhibition of OXA-57 by selected β-lactams and β-lactamase inhibitors.

| Time (min) | Inhibitor | pIC<sub>50</sub> | IC<sub>50</sub> (μM) |
| --- | --- | --- | --- |
| 10 | Clavulanate | 4.5 ± 0.03 | 30.3 |
| | Avibactam | 4.2 ± 0.03 | 58.8 |
|  | Vaborbactam | <2.4 | - |
|  | Aztreonam | <2.4 | - |
|  | Cefotaxime | <2.4 | - |
|  | Ceftazidime | <2.4 | - |
| 30 | Clavulanate | 4.9 ± 0.04 | 12.3 |
| | Avibactam | 4.7 ± 0.04 | 18.8 |
|  | Vaborbactam | <2.4 | - |
|  | Aztreonam | <2.4 | - |
|  | Cefotaxime | <2.4 | - |
|  | Ceftazidime | <2.4 | - |
| 60 | Clavulanate | 5.0 ± 0.04 | 9.4 |
| | Avibactam | 4.9 ± 0.04 | 13.2 |
|  | Vaborbactam | <2.4 | - |
|  | Aztreonam | <2.4 | - |
|  | Cefotaxime | <2.4 | - |
|  | Ceftazidime | <2.4 | - |

As well as being effective antibiotics, some β-lactams efficiently inhibit certain β- lactamases. As steady-state measurements (above) failed to detect OXA-57-catalyzed hydrolysis of the 3^rd^ generation (oxyimino)cephalosporins cefotaxime or ceftazidime, or the monobactam aztreonam, we investigated their ability to inhibit OXA-57 via formation of long-lived acyl-enzyme species. However, we could not determine IC_50_ values for any of these compounds from measurements at concentrations up to 4 mM (**Table 2**), indicating that these β-lactams do not form stable complexes at the OXA-57 active site.

### Crystal Structure of Uncomplexed OXA-57

While some biochemical characterization of OXA-57 has been reported, no crystal structure has been available. Accordingly, purified OXA-57 was crystallized from two different conditions in an identical crystal form (space group *C*2221 with one molecule in the asymmetric unit) and the structure solved to resolutions of 1.8 and 2.0 Å, respectively (**Table S4**). The two structures are effectively identical (RMSD 0.31 Å as determined by SSMSuperpose^36^), although in neither case could we model a continuous polypeptide chain, with residues 209 -214 (comprising the β4 - β5 loop adjacent to the active site) and 243 – 245; and 209 – 214 and 243 – 248; omitted from the final models derived from the 2.0 Å and 1.8 Å datasets, respectively, due to poorly defined electron density in these regions. The discussion below, and associated molecular simulations, are based upon the 2.0 Å resolution structure as the model is more complete, although the data are of slightly lower quality.

The final modelled structure consists of eight α-helices surrounding a six-stranded antiparallel β-sheet (**Figure 1a**) with the conserved SXXK motif containing the nucleophilic serine residue^29^ at the N-terminus of the α1 helix (residues S53 and K56). Of the two further sequence motifs conserved in OXA β-lactamases the SXV motif, whose serine (OXA-57 S104) relays protons in the class-D β-lactamase reaction mechanism^14,22^ (**Figure S1**), sits on the loop between helices α3 and α4 and the KTG motif (OXA-57 residues 201 – 203) on the β4 strand. The Ω-loop (OXA-57 residues 137 - 152), a flexible region important to catalytic activity and substrate selectivity in class A β-lactamases^38^, is well defined by the experimental electron density and could be modelled in its entirety. Overall, the fold of OXA-57 is similar to those of other class-D β-lactamases (root-mean-square deviation values [RMSDs] for OXA-57 compared with selected OXA structures are presented in **Table S5**). Electron density was also well defined for mechanistically important residues in the OXA-57 active site; inspection of difference maps clearly showed that, consistent with previous studies^14^, K56 is fully carbamylated (**Figure 1b**, in both uncomplexed structures refined with occupancy 1.0 and B-factors for carbamate atoms equivalent to those of the K56 Nε atom (**Table S6**)), and forms hydrogen bonds with S53 and with W149 on the Ω-loop.

**Figure 1.**
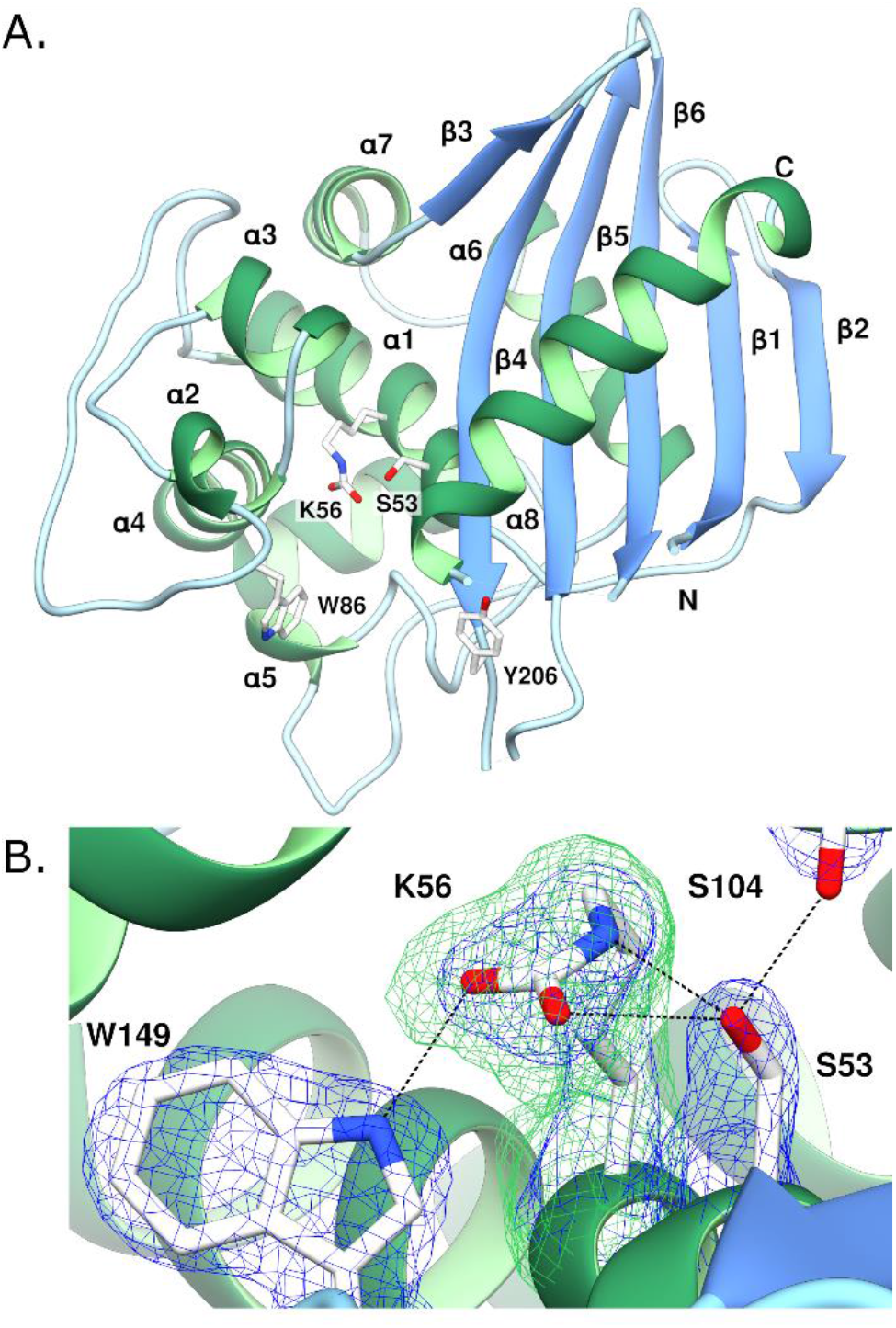
Crystal Structure View of OXA-57. **A.** Ribbon diagram of the overall structure, showing secondary structure elements and key residues. **B.** Active site showing 2*F*_o_ - *F*_c_ electron density (blue mesh, 1.5σ) for residues S53, K56, S104 and W149. Omit *F*o - *F*c map (green) calculated with K56 removed and contoured at 3σ shows carbamylation of K56. Hydrogen bonds are shown as dashed lines. The figure was generated using UCSF Chimera^37^.

### Structural Basis of OXA-57 by Inhibition by Avibactam

To assess the effect of inhibitor exposure on the structure of OXA-57, crystals were soaked with avibactam for 16 hours. Diffraction data at 2.48 Å resolution (**Table S4**) showed clear *F*_o_- *F*_c_ density in the active site that could confidently be modelled as ring-opened avibactam covalently attached to S53 (real-space correlation coefficient (RSCC) 0.96, **Figure 2a, b**). Soaking with avibactam did not alter the space group (*C*222_1_) or unit cell dimensions in comparison to those for the uncomplexed enzyme (**Table S4**) and avibactam induces no substantial changes to the overall structure (C_α_ RMSD 0.31 Å between the native and avibactam-bound structures). Avibactam was refined at full occupancy with B-factors comparable to those for protein atoms (**Table S4**).

**Figure 2.**
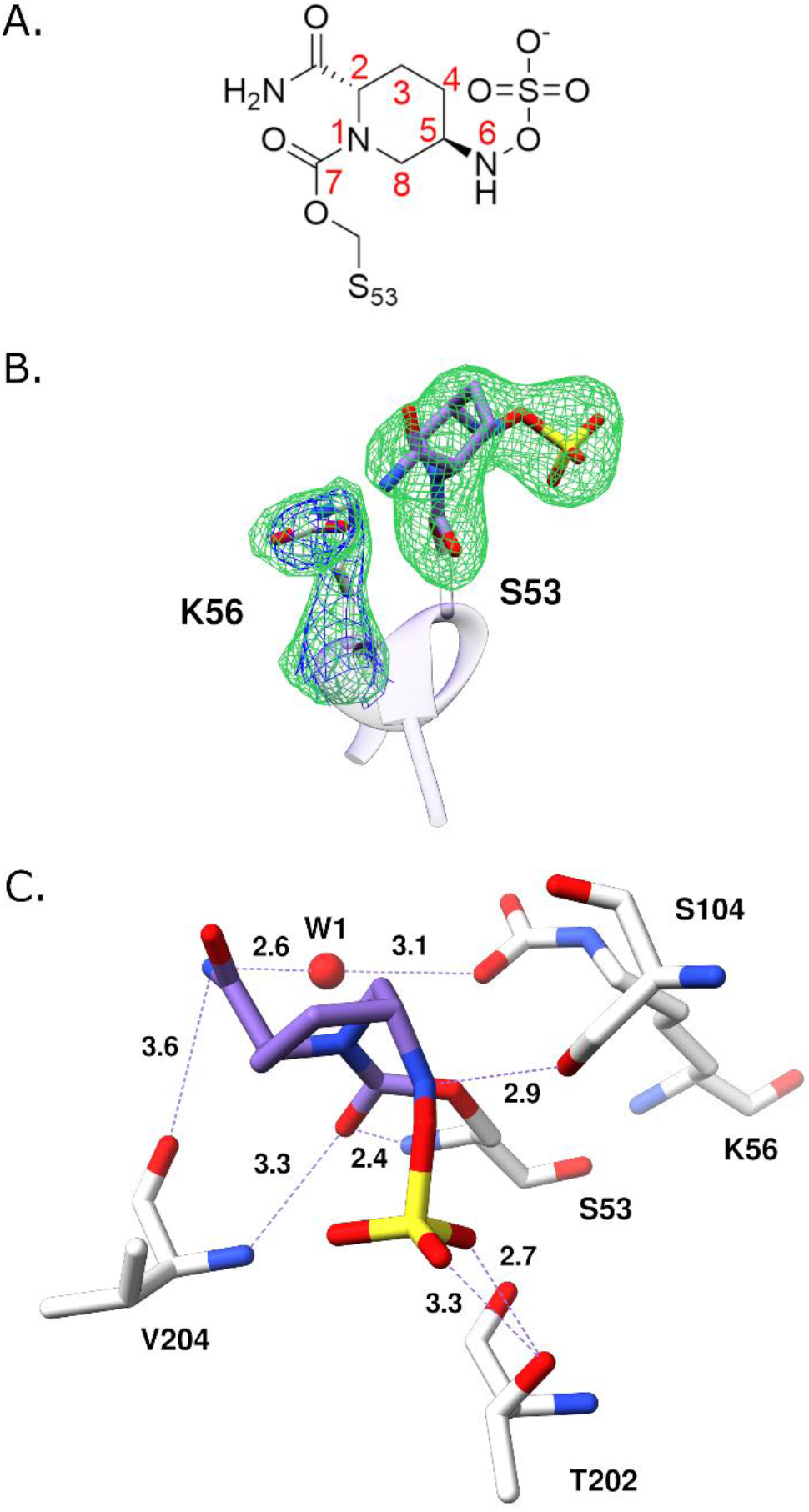
OXA-57:Avibactam Complex. **A.** Schematic of avibactam:enzyme complex showing avibactam atom numbering. **B.** Electron density of avibactam bound to OXA-57. 2*F*_o_

The opened six-membered piperidine ring of avibactam adopts a chair-like conformation, as seen in other SBL:avibactam complexes^39,40^, with binding stabilized by an extensive network of hydrogen bonds (**Figure 2c**). The avibactam C7 carbonyl oxygen interacts with the - *F*_c_ electron density (blue, 1.5σ) shows carbamylation of K56. Green density shows Omit *F*_o_ - *F*_c_ map (3σ) calculated with avibactam and carbamylated K56 removed. **C.** Interactions of avibactam with the OXA-57 active site. Avibactam carbon atoms are colored purple, non-carbon atoms by atom type, interacting residues are shown as white sticks, and water molecules as red spheres. Hydrogen bonding interactions are shown as dashed lines with distances between heavy atoms (Å) shown. This Figure was generated using ChemDraw and UCSF Chimera^37^. backbone amides of S53 (2.4 Å) and V204 (3.3 Å) on the neighboring β4 strand. The avibactam-derived C2 carboxamide hydrogen bonds with the backbone carbonyl of V204 (3.6 Å). The avibactam-derived sulfate is stabilized by interactions between two of its oxygen atoms and the side chain hydroxyl of T202 (2.7 Å and 3.3 Å, respectively). In contrast to some complexes with other serine β-lactamases^41–43^, our structure provides no evidence for desulfation of bound avibactam. K56 remains fully carbamylated (**Figure 2b)** as evidenced by B-factors for the carbamate group (**Table S6**); the carbamate is reoriented compared to its position in the uncomplexed structure (**Figure S4**), enabling hydrogen bonding to the avibactam C2 carboxamide via a water molecule (**Figure 2c** W1, 2.6 and 3.1 Å). The urea-derived N6 nitrogen of avibactam is orientated towards the carbonyl group of the acyl-enzyme-type complex, apparently stereo-electronically favoring reaction to reform avibactam, consistent with the reversible nature of avibactam inhibition^44^. In prior structures, the N6 nitrogen of avibactam and related diazabicyclooctanes has been observed both in this conformation and one which is orientated away from the carbonyl group of the acyl-enzyme type complex^45,46^.

### Interactions of OXA-57 with hydrolyzed meropenem

To investigate the structural basis of OXA-57-catalyzed β-lactam (specifically carbapenem) hydrolysis, OXA-57 crystals were soaked with meropenem for 1 hr before freezing for subsequent diffraction experiments. Diffraction data extending to 2.42 Å resolution (**Table S4**) revealed positive *F*_o_-*F*_c_ density in the active site consistent with ring-opened meropenem covalently linked to S53. Meropenem exposure did not affect the space group (*C*2221) or unit cell dimensions compared to those for the native enzyme (**Table S4**) and induced minimal changes to the overall structure, (Cα RMSD 0.38 Å).

Electron density was well defined for the meropenem derived pyrroline ring, carbonyl oxygen, C2 exocyclic sulfur and C6 hydroxyethyl group (**Figure 3b**), but was weak for the remainder of the C2 substituent and C-3 carboxylate. At this resolution the orientation of the 6-α-hydroxyethyl group could not be unambiguously assigned, this was therefore modelled in an orientation, pointing away from the carbamylated K56, similar to that observed in carbapenem complexes of other class D β-lactamases^47–50^. The modelled ligand has an RSCC of 0.9 as defined in Phenix. Carbapenem hydrolysis by serine β-lactamases can generate alternative products; tautomerization of the Δ2-enamine product (in which the carbapenem C2 is *sp^2^* hybridized) results in the Δ1-imine in two equilibrating isomeric forms (with C2 *sp*^3^ hybridized)^16^ (**Figure S1**). Based upon experimental electron density maps, which indicated the exocyclic C2 sulfur to be coplanar with the pyrroline ring, the meropenem-derived acyl-enzyme was modelled as the Δ2 enamine, though the presence of other tautomers is possible (**Figure 3A, B**).

**Figure 3.**
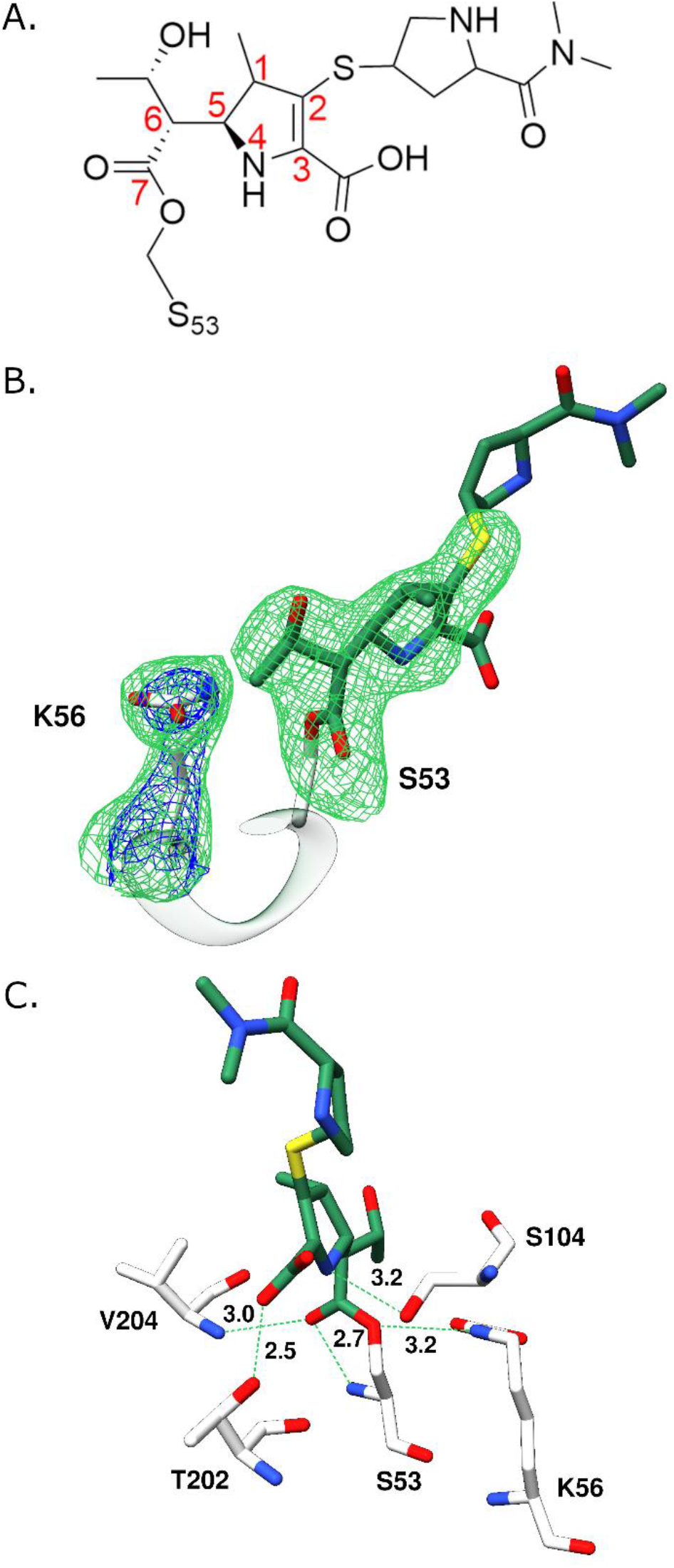
The Meropenem Acyl-enzyme of OXA-57. **A.** Schematic of the meropenem acyl-enzyme in the Δ^2^ enamine tautomer. **B.** Electron density for meropenem bound in the active site of OXA-57. An Omit *F*o - *F*c map (green, 3σ) was calculated with meropenem and the carbamylated K56 removed. **C.** Key interactions of the OXA-57:meropenem acyl-enzyme. Meropenem carbon atoms are colored green, all other non-carbon atoms colored by atom type. Interacting residues are white sticks, and hydrogen bonding interactions are shown as dashed lines with distances between heavy atoms labelled in Å. This Figure was generated using ChemDraw and UCSF Chimera^37^.

A network of hydrogen bonds stabilizes the meropenem derived acyl-enzyme (**Figure 3C**). The S53 oxygen atom interacts with the side chain amide of K56 (3.2 Å), which *F*o-*F*c electron density maps clearly reveal to be fully carbamylated and for which the carbamate was refined at full occupancy (**Table S6**). The acyl-enzyme carbonyl oxygen is positioned, *via* hydrogen bonds, in the oxyanion hole formed by the backbone amides of S53 and V204 (2.7 Å and 3.0 Å). The pyrroline nitrogen N4 hydrogen bonds to the S104 hydroxyl (3.2 Å) and the C3 carboxylate interacts with the side chain O of T202 (2.5 Å). However, no electron density was evident for an active site water molecule positioned for deacylation of the meropenem acyl-enzyme.

### Molecular Dynamics Simulations of Native OXA-57 and the OXA-57:Meropenem Acyl-enzyme Complex

To investigate the dynamic behavior of OXA-57, and the effects upon it of acylation by meropenem, we ran five 500 ns molecular dynamics (MD) simulations (repeats 1 - 5, R1 - R5) of uncomplexed OXA-57 and the OXA-57:meropenem complex (totaling 2.5 µs per system). Our crystal structures were used as starting points for the simulations, with unstructured regions (corresponding to the β4 - β5 and β6 - α8 loops in native, uncomplexed OXA-57; and loops α1 - α2, β4 - β5 and β6 - α8 in the OXA-57:meropenem acyl-enzyme complex) constructed as described in Methods (**Figure S5**). The time dependence of backbone RMSD values during each simulation is shown in **Figures S6** and **S7**. In some cases, we observed variations in backbone RMSD over the course of the 500 ns simulations; these were subsequently recalculated excluding regions corresponding to the β4 - β5 loop (residues S205 - S220) and the C-terminus (residues L266 - R269) resulting in RMSDs that stabilized around ∼ 1.0 Å (**Figures S8, S9**). These averaged 0.9, 1.0, 0.9, 1.0, and 1.3 Å for simulations R1 - 5 of native OXA-57; and 1.1, 1.0, 1.0, 1.1, and 1.0 Å for simulations of the OXA-57:meropenem complex. There were no marked differences in residue-based backbone RMSF distributions between the simulations of native OXA-57 and the meropenem acyl-enzyme (**Figure S10**). In both cases, the β4 - β5 loop is clearly the most mobile region of the protein. For the acyl-enzyme, RMSF values in the region corresponding to the β4 - β5 loop increased in one replicate simulation (R3), although this was not evident in the other four simulations. Snapshots of structures taken at 100 ns intervals throughout the MD trajectories of uncomplexed OXA-57 and the meropenem acyl-enzyme indicate that the β4 - β5 loop accesses similar conformations, and undergoes similar motions, regardless of the presence of meropenem (**Figures S11** and **S12**).

The carbamylated lysine sidechain (residue K57) is stable over the course of the simulations of both native OXA-57 and the OXA-57:meropenem acyl-enzyme (average RMSDs 0.5 and 0.6 Å, respectively (**Figures S13** and **S14**)). However, in the meropenem acyl-enzyme, bound meropenem deviates from the crystallographically observed conformation within the first 100 ns of each simulation, with movements of the exocyclic sulfur and pyrrolidine ring of the C2 substituent largely accounting for the increase in RMSD. Bound meropenem adopts different conformations throughout the simulations, ultimately favoring a conformation approximately 5 - 6 Å distant from that observed in the crystal structure (**Figure S16**). During the MD simulations the C2 substituent (R-group) usually resides between the α1 - α2 and β4 - β5 loops (**Figure S16**), rather than pointing into bulk solvent.

Hydrogen bonding analysis indicates that bound meropenem forms five hydrogen bonds to OXA-57 that are present in at least three of the five repeat simulations and over more than half of the MD trajectory (**Figure 5**). The meropenem carbonyl oxygen (O25) is positioned in the oxyanion hole formed by the backbone amides of V204 (average hydrogen bond occupancy 95% over the five MD simulations), and S53 (average occupancy 97%). The acyl-enzyme ester oxygen (OG) is hydrogen bonded to the side chain N01 of carbamylated K56 (72% occupancy). The oxygen atom (O22) of the meropenem C6 α-hydroxyethyl group is hydrogen bonded to atom OQ2 of the carbamylated lysine K56 for on average 84% of the simulation time and is thus orientated towards K56 for the majority of the MD trajectory. A fifth hydrogen bond, between N15 in the pyrrolidine ring of the meropenem C2 substituent and the backbone amide nitrogen of G88 located in the α1 - α2 loop, is evident for over half of the trajectory in 4/5 of the repeat MD simulations (excepting R4). This interaction likely helps stabilize the “upwards” conformation of the portion of the meropenem C2 substituent that lies beyond the exocyclic sulfur (**Figure 5a**).

**Figure 4.**
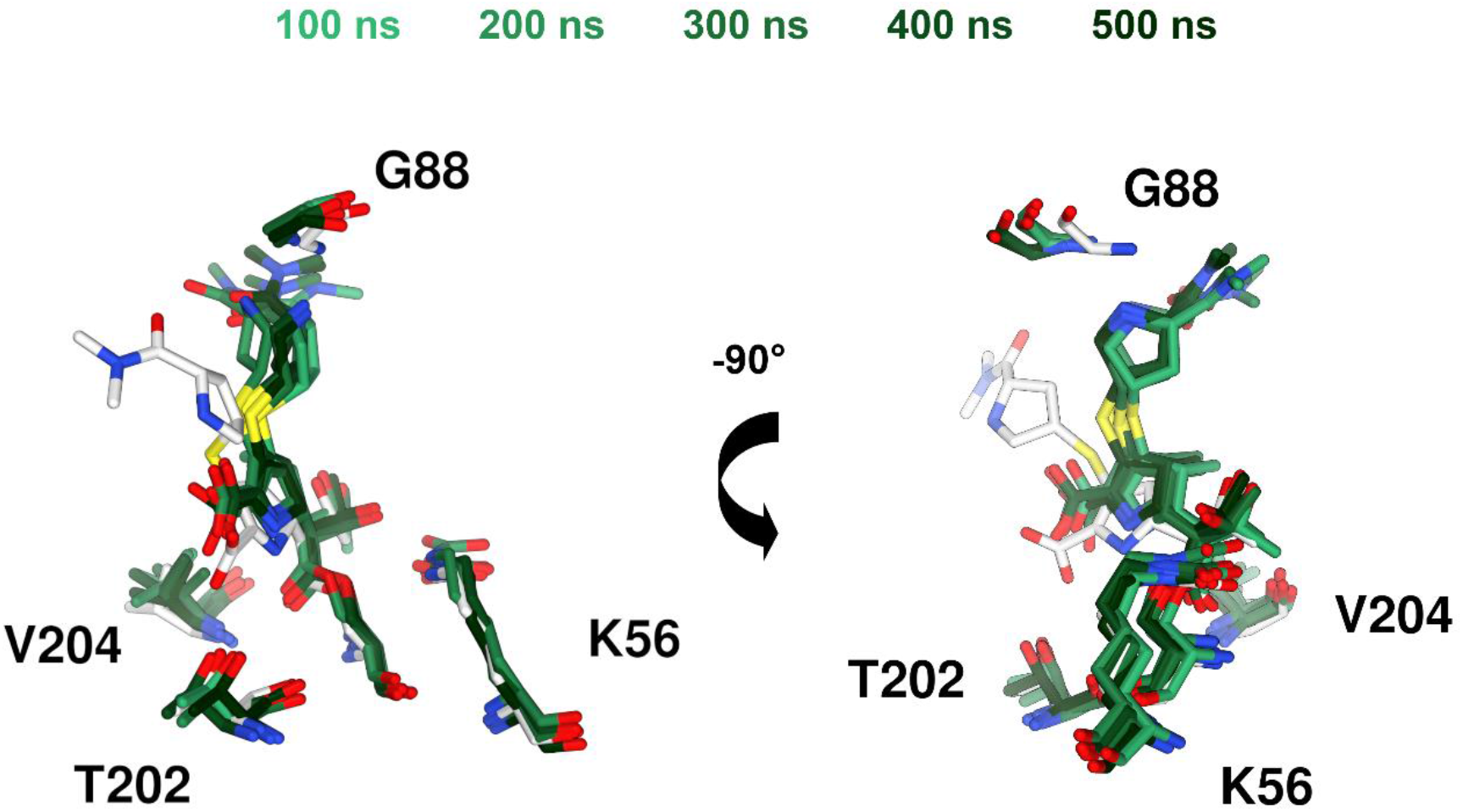
Relocation of Bound Meropenem in the OXA-57 Active Site. Superpositions of bound meropenem in snapshots taken at 100 ns intervals throughout the trajectory of MD simulations of the OXA-57:meropenem acyl-enzyme. Data shown are from simulation R5. (Additional data are provided in Supporting Information (**Figure S16**)). Carbon atoms in the crystal structure are in white; carbon atoms from simulations are in green, snapshots at successive 100 ns intervals are shaded incrementally from light (100 ns) to dark (500 ns) green. Other atom colors as standard. This Figure was generated using UCSF Chimera^37^.

**Figure 5.**
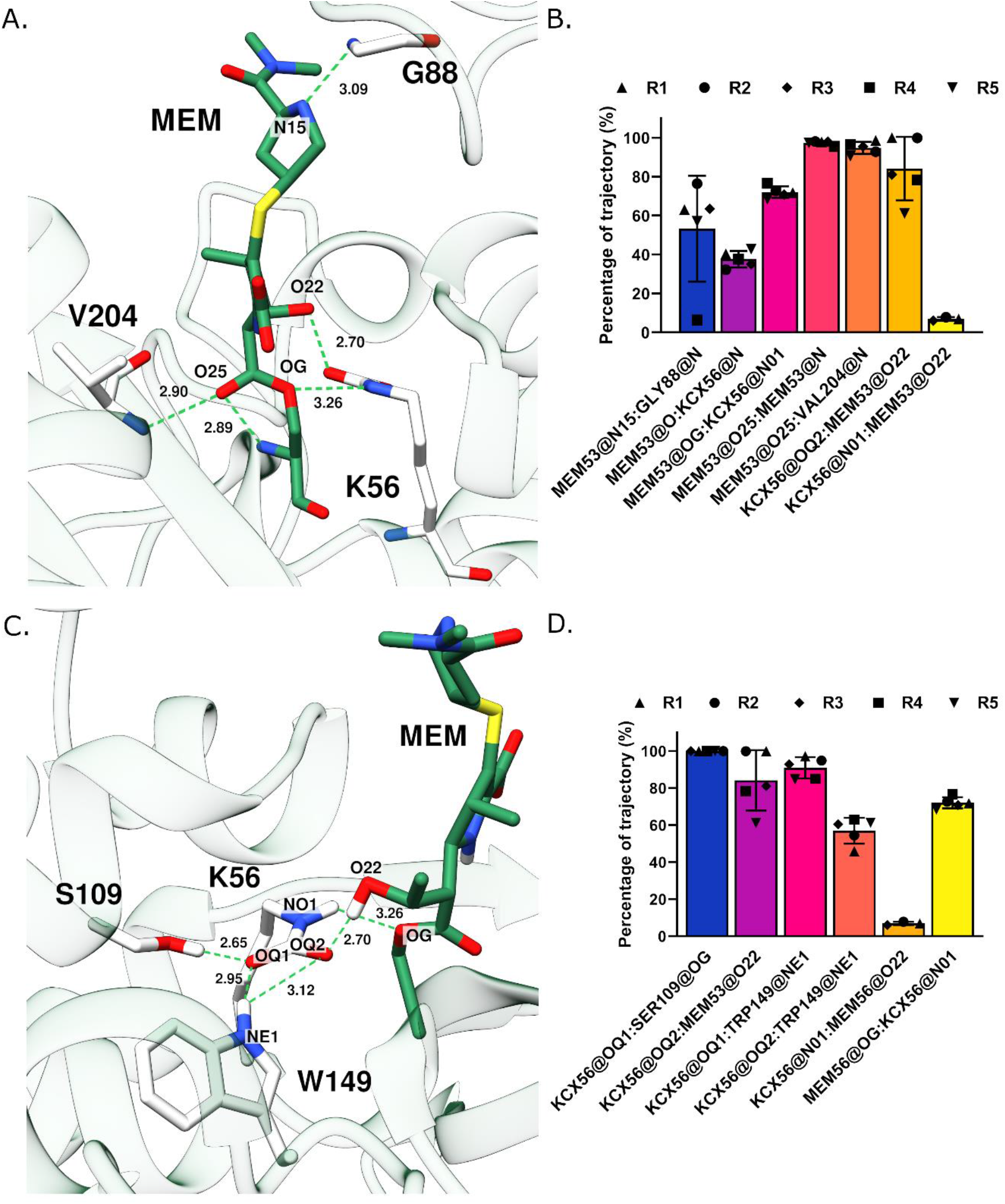
Hydrogen Bonding Networks Between Meropenem and OXA-57. **A.** Hydrogen bond interactions between OXA-57 and meropenem mapped onto a representative structure from simulations of the OXA-57:meropenem acyl-enzyme. **B.** Hydrogen bond interactions between OXA-57 and meropenem expressed as the mean percentage occupancy during 425 ns trajectories of simulations of the OXA-57:meropenem acyl-enzymes. **C.** Hydrogen bond interactions between carbamylated K56 and meropenem mapped onto a representative structure from simulations of the OXA-57:meropenem complex. **D.** Hydrogen bond interactions between carbamylated K56 and meropenem expressed as the mean percentage occupancy during 425 ns trajectories of the OXA-57:meropenem complex simulations. For **A** and **C**, the representative structure has the lowest RMSD to the average coordinates of all five repeat simulations of the OXA-57:meropenem acyl-enzyme. For clarity, only hydrogen bonds present for over half of the trajectories in at least three simulations are shown. Hydrogen bond distances shown are averages over all five simulations. For panels **B** and **D**, bar heights denote mean values, with error bars representing standard deviations. Values for individual repeats are shown as upward triangles (R1), circles (R2), diamonds (R3), squares (R4) and downward triangles (R5). Atoms involved in hydrogen bonds are specified as acceptor:donor. For clarity, only hydrogen bonds present in at least three simulations are shown. Meropenem (MEM) carbon atoms are shown in green, protein carbon atoms in white, other atom types atoms (including hydrogen atoms where involved in hydrogen bonds) in standard colors. Figure was generated using UCSF Chimera^37^ and GraphPad Prism.

The crystal structure of the OXA-57:meropenem acyl-enzyme suggests the presence of a hydrogen bond between the meropenem C3 carboxylate and the side chain hydroxyl of T202, although experimental electron density does not well define the orientation of the C3 carboxylate group. However, such an interaction is rarely evident during the MD simulations (**Figure S17**). Instead, the meropenem C3 carboxylate is occasionally hydrogen bonded to S104 (in on average approximately 4% of the trajectories via atom O01, and 6% via O03) but, for most of the simulations, faces the bulk solvent and is coordinated by 1 - 6 (most commonly 3) water molecules (**Figure S17**). During the MD simulations we observe that the meropenem C3 carboxylate can form water-mediated interactions with the T202 hydroxyl group, and additionally with the hydroxyl group of T202 and atom H01 attached to the nitrogen of the meropenem pyrroline ring (**Table S7**). The high degree of mobility of, and absence of persistent interactions made by, the meropenem C3 carboxylate throughout the MD simulations is consistent with the poor electron density observed for this group in the crystal structure.

We also examined the hydrogen bonding network made by the carbamylated lysine K56 (**Figure 5c**). In addition to the hydrogen bond to meropenem O22 (the C6 α-hydroxyethyl group, 84%, above), the K56 carbamyl group makes a further two hydrogen bonding interactions: of OQ1 to the side chain hydroxyl of S109 (100% average occupancy), and of either OQ1 or OQ2 to the indole N atom NE1 of W149 (91/57%; **Figure 5d**). Atom N01 of the carbamylated lysine sidechain makes an additional hydrogen bond to the meropenem C6 α- hydroxyethyl oxygen O22 (> 60% occupancy in all five trajectories). Involvement in multiple high-occupancy hydrogen bonds ensures that carbamylated K56 retains a tightly defined conformation throughout the MD trajectories.

Resolution of β-lactam acyl-enzymes of SBLs occurs through addition of water to the acyl- enzyme carbonyl, releasing the hydrolyzed product and regenerating active enzyme. However, our crystal structure of the OXA-57:meropenem acyl-enzyme did not contain any water molecules appropriately positioned for deacylation. Accordingly, the MD trajectories were investigated to search the OXA-57 active site for candidate deacylating water molecules. Water molecules within 10 Å of both the acyl-enzyme carbonyl carbon (meropenem C24) and OQ2 of carbamylated lysine K56 (the most likely general base for activation of an incoming water molecule, **Figure 6A**) were extracted from each frame of each of the five MD trajectories (42, 501 frames in total) and their positions and occupancies plotted (**Figure 6B**).

**Figure 6.**
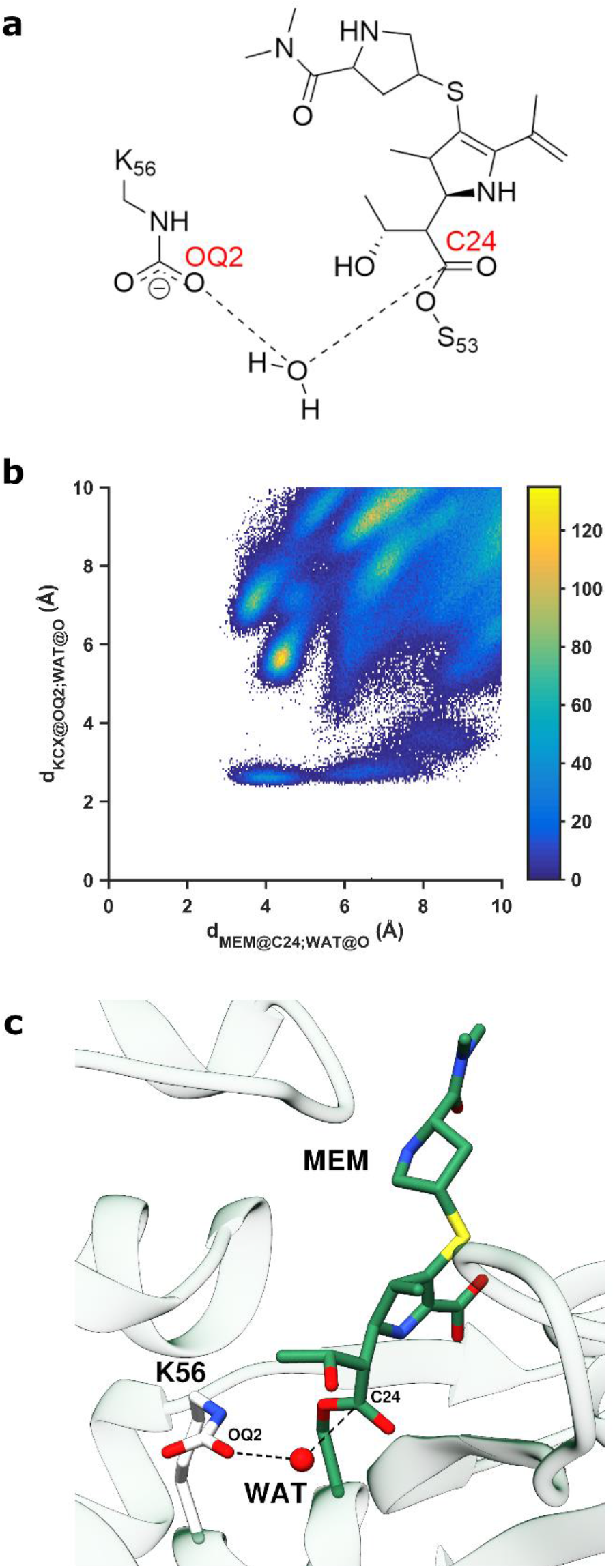
Candidate Deacylating Water Molecules are Transiently Present During MD Simulations of the OXA-57:Meropenem Acyl-enzyme. **A**. Active site of the OXA- 57:meropenem acyl-enzyme. Distance distributions calculated in **B** are shown as dashed lines. **B**. Distance distributions of water molecules between meropenem C24 (carbonyl carbon) and K56 OQ2 over 425 ns MD trajectories of the OXA-57:meropenem acyl-enzyme. Data shown are from simulation R4. (Data for repeat simulations R1 - 3 and R5 can be found in Supporting Information (**Figure S18**)). **C**. Hydrogen bond interactions between K56, meropenem and water in the deacylating position mapped onto a representative structure from simulation R4 of the OXA-57:meropenem acyl-enzyme. Figure was generated using ChemDraw, MATLAB and UCSF Chimera.^37^

In four of the five repeat simulations water molecules positioned for deacylation were either entirely absent (repeats R1 and R3) or present to a negligible degree (repeats R2 and R5, **Figure S18**). In simulation R4, some water molecules were observed located within 3.5 Å of both meropenem C24 and K56 OQ2, and thus favorably positioned for deacylation (**Figure 6B**), although in most frames water molecules are between 4 – 5 Å distant from meropenem C24 and 5 – 6 Å from K56 OQ2. In simulations R2, R4 and R5, water molecules reside within 3.5 Å of both meropenem C24 and K56 in a total of 1, 602, and 32 frames, equating to 0.002, 1.4, and 0.08% of the simulations, respectively. Water molecules occupying this potentially deacylation-competent position can exchange with bulk solvent; in simulation R4, 3 individual water molecules are observed in this position, while 9 are observed in R5. Furthermore, water molecules positioned for possible deacylation also interact with the meropenem 6-α-hydroxyethyl group. Specifically, meropenem atom O22 (the hydroxyl of the C6a- hydroxyethyl substituent) lies within 3.5 Å of the candidate deacylating water, when present, in 601/602 and 25/32 of the frames in repeat simulations R4 and R5, respectively (**Figure 6c**). This finding suggests that the hydrogen bond between meropenem O22 and K56 OQ2 must break for a water molecule to occupy the deacylating position. In simulation R4, where candidate deacylating water molecules were most prevalent, meropenem O22 is not hydrogen bonded to K56 OQ2 when the water molecules are present. In simulations R2 and R5 there are very rare instances (one frame in R2, five in R5) when both a water molecule and the meropenem O22: K56 OQ2 hydrogen bond are observed. Taken together, however, these analyses indicate that the active site of the OXA-57:meropenem acyl-enzyme does not readily accommodate a water molecule positioned for deacylation, and that the existence of a hydrogen bond between meropenem O22 (the hydroxyethyl group) and the carboxylate of carbamylated K56 may be a contributing factor to this.

Two further factors have been identified as contributors to deacylation and carbapenem turnover in OXA β-lactamases. First, it has been suggested that the positioning and movement of a leucine residue (L166 in OXA-23) may play a critical role in permitting access of a water molecule to the active site to enable deacylation^50^. In OXA-57 the analogous residue is an isoleucine (I150) located on the α5-α6 loop. Accordingly, we interrogated our simulations seeking correlation between motions of I150 and the presence/absence of a water molecule positioned for deacylation, but none was evident. Secondly, in some CHDLs, a hydrophobic bridge flanking the active site is either present or forms during carbapenem hydrolysis^51,52^, with the residues involved varying between OXA family members. Inspection of the OXA-57 structure identifies residues W86 and Y206, on the α1-α2 and β4-β5 loops, respectively (**Figure 1A**), as candidates to form a hydrophobic bridge across the active site cleft during the hydrolysis reaction, although they do not interact in either the uncomplexed or meropenem acyl-enzyme crystal structures. We calculated the minimum distance between any two carbon atoms in these residues during the 500 ns MD simulations of both the uncomplexed and meropenem acyl-enzyme structures and observed that they rarely approached within ≤ 4.0 Å of one another. Hence our simulations do not support a role for I150 in regulating access of water to the OXA-57 active site for deacylation, and do not identify any propensity for formation of a hydrophobic bridge across the OXA-57 active site cleft.

## 3. Discussion

Our kinetic studies, in agreement with previous work^22^, show OXA-57 to be a penicillinase that also hydrolyzes 1^st^ generation cephalosporins (cepahalothin) and which is inhibited by clavulanate at micromolar concentrations. Further, we show OXA-57 to be a weak carbapenemase with kinetic parameters for carbapenems comparable to those previously reported for OXA-10 (**Table S2**). Previous work indicates that class-D β-lactamases with apparent narrow-spectrum activity can also hydrolyze carbapenems when expressed in their native hosts, posing a significant clinical risk^53^. Despite the weak activity of OXA-57 towards carbapenems, it is possible that its over-expression could result in clinically relevant carbapenem MICs, as previously seen for the *B. pseudomallei* class A β-lactamase PenI^8,54^. Significantly, efflux has also been reported as a contributing factor to carbapenem resistance in other Gram-negative pathogens^55,56^. *B. pseudomallei* has 3 characterized resistance nodulation division (RND) efflux pumps which are utilized in resistance to other clinically important antibiotics^57^. It is therefore possible that the weak carbapenemase activity of OXA- 57, when present in a relatively impermeable host organism such as *B. pseudomallei*, and potentially in combination with upregulated efflux systems, may contribute to imipenem or meropenem resistance phenotypes in *B. pseudomallei* clinical isolates.

Of the tested compounds, clavulanate was the most effective β-lactamase inhibitor against OXA-57. In comparison with other class-D β-lactamases, avibactam was up to two orders of magnitude less active towards OXA-57, while OXA-57 showed very limited susceptibility towards vaborbactam, consistent with its weak activity towards other class D enzymes (**Table S3**). Avibactam, like other DBO inhibitors, functions via a reversibly binding mechanism where a covalent inhibitory complex formed by carbamylation of the nucleophilic serine can undergo slow decarbamylation and recyclization that regenerates active inhibitor^58^. For some SBLs, such as KPC-2 (class A) and AmpC (class C)^41–43^, avibactam and other DBOs can undergo desulfation leading to inhibitor degradation/hydrolysis and release of inactive products^41,59^. Our crystal structure of the OXA-57:avibactam complex (obtained from crystals exposed to avibactam for 16 h before data collection), provides no evidence of desulfation, indicating that, as observed for other OXA enzymes^39,41^, OXA-57 is likely to support reversible decarbamylation of DBOs through recyclization.

Avibactam binds the OXA-57 active site in a similar orientation to that observed in complexes with other class-D β-lactamases (OXA-10, OXA-24, OXA-48; **Figure 7**). In these four OXA:avibactam complex structures, one avibactam sulfate oxygen makes a hydrogen bond to a neighboring threonine (OXA-57 T202). In the OXA-57 complex this is the only hydrogen bond made by the avibactam sulfate, but in comparator structures the sulfate group makes further hydrogen bonds to a conserved arginine residue (**Figure 7**). OXA-57 contains an equivalent arginine (R255), but this is unfavorably oriented for interaction with the avibactam sulfate. Both OXA-10 and OXA-24 make additional interactions with avibactam that are absent from the OXA-57 complex; namely an additional hydrogen bond between the avibactam sulfate oxygens and the serine residue of the SXV motif (OXA-10 S115, OXA-24 S128; **Figure 7a, b**). In the OXA-24 complex, the avibactam sulfate further interacts with two active site water molecules (**Figure 7b**). The limited extent of interactions made with avibactam are consistent with its reduced potency towards OXA-57, compared to other class D enzymes.

**Figure 7.**
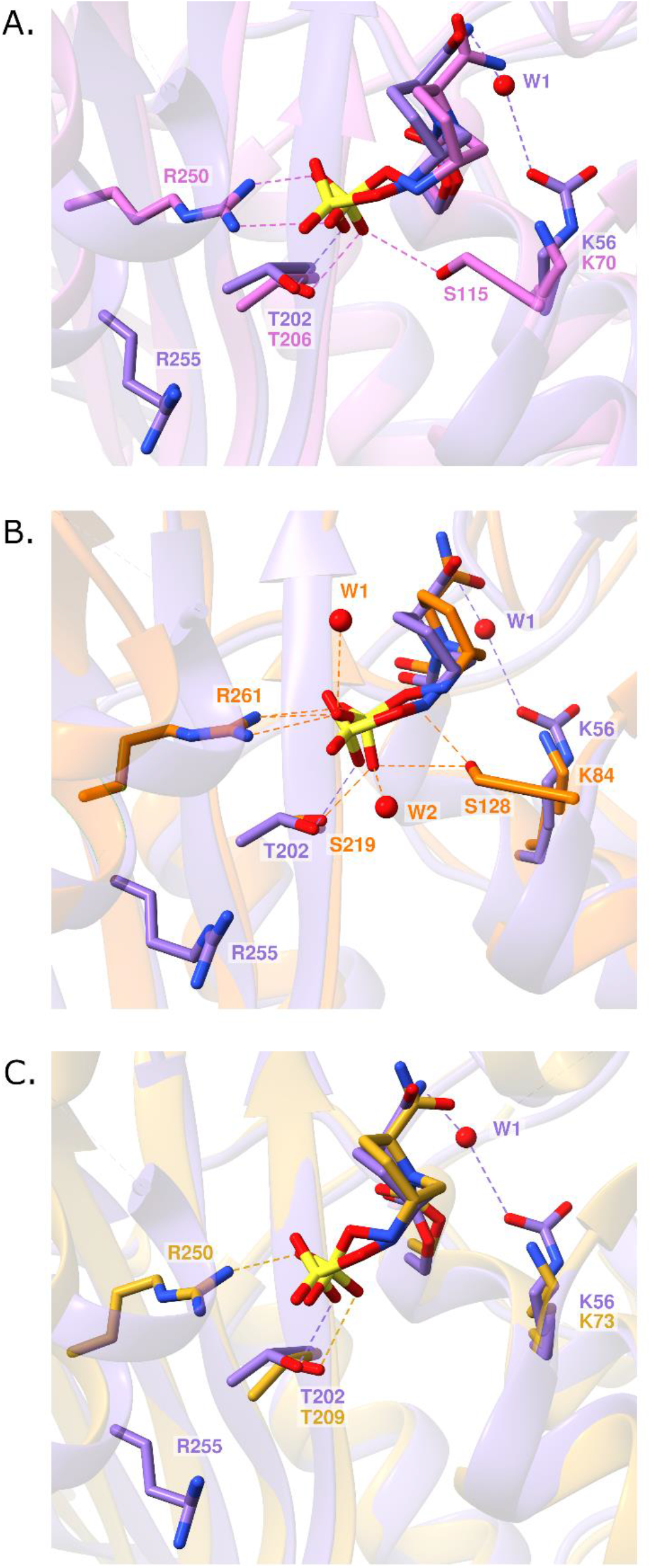
Avibactam Binding to Class D β-lactamases. **a.** Overlay of avibactam binding to OXA-57 (purple) and OXA-10:avibactam complex (PDB:4S2O^39^, pink). **b.** Overlay of avibactam binding to OXA-57 (purple) and OXA-24:avibactam complex (PDB:4WM9^40^, orange). **c.** Overlay of avibactam binding to OXA-57 (purple) with OXA-48:avibactam complex (PDB:4WMC^40^, gold). Non-carbon atoms are colored by atom type, hydrogen bonding interactions are shown as dashed lines. RMSD values between structures can be found in Supporting Information (**Table S8**). The Figure was generated using UCSF Chimera^37^.

The most notable difference between the OXA:avibactam complexes is the presence of the carbamylated lysine in OXA-57. Carbamylation of the OXA active site lysine is pH-dependent, and favored under basic conditions^14^, as utilized in crystallization of OXA-57 (OXA-10, pH 6.5; OXA-24, pH 6.0; OXA-48, pH 7.5 and OXA-57, pH 8.5). Although carbamylation of the active site lysine has been suggested to be disfavored in avibactam complexes of OXA β- lactamases^39^, in the OXA-57:avibactam complex the carbamylated lysine makes a water- mediated interaction with the avibactam C2 carboxamide (**Figure 7**), potentially stabilizing both the lysine carbamate and bound avibactam. The role of lysine carbamylation in susceptibility to inhibition of OXA enzymes by avibactam (and other DBOs) however remains to be clarified.

In comparison to other CHDLs, *K*M values for hydrolysis of the carbapenems imipenem and meropenem by OXA-57 are in most cases 5 – 10-fold higher, and *kcat* values an order of magnitude lower (**Table S2**). Such differences may reflect factors including altered hydrogen bonding networks with bound substrates, and differences in active site accessibility to the water molecule required for deacylation. Superimposition of our OXA-57:meropenem structure on those of other OXA:carbapenem acyl-enzyme complexes identifies differing interactions with bound carbapenems (**Figure 8**). Hydrogen bonds in common with the OXA-1:doripenem and OXA-10:imipenem complexes (**Figure 8a, b**), which also contain a carbamylated active site lysine, include those between the carbapenem acyl-enzyme carbonyl oxygen and the oxyanion hole formed by the backbone amide nitrogens of the serine nucleophile and the residue immediately following the conserved KTG motif (OXA-57 V204, OXA-1 A215, OXA-23 W219, and OXA-48 Y211; **Figure 8a, c, d**). Similarly, a hydrogen bond between the carbapenem C3 carboxylate and the threonine residue of the active site KTG motif (OXA-57 T202) is present in all of these complexes. Interestingly, this hydrogen bond did not persist during MD simulations, which identified bound meropenem as relatively mobile within the OXA-57 active site. It is also important to note that, due to weak experimental electron density, we were unable to definitively assign the orientation of the carbapenem 6-α-hydroxyethyl group in the OXA-57:meropenem acyl-enzyme complex crystal structure, and it was therefore modelled in an orientation, close to the carbamylated K56, based on that observed in other OXA:carbapenem complexes. This interaction persisted during molecular dynamics simulations of the OXA-57:meropenem acyl-enzyme.

**Figure 8.**
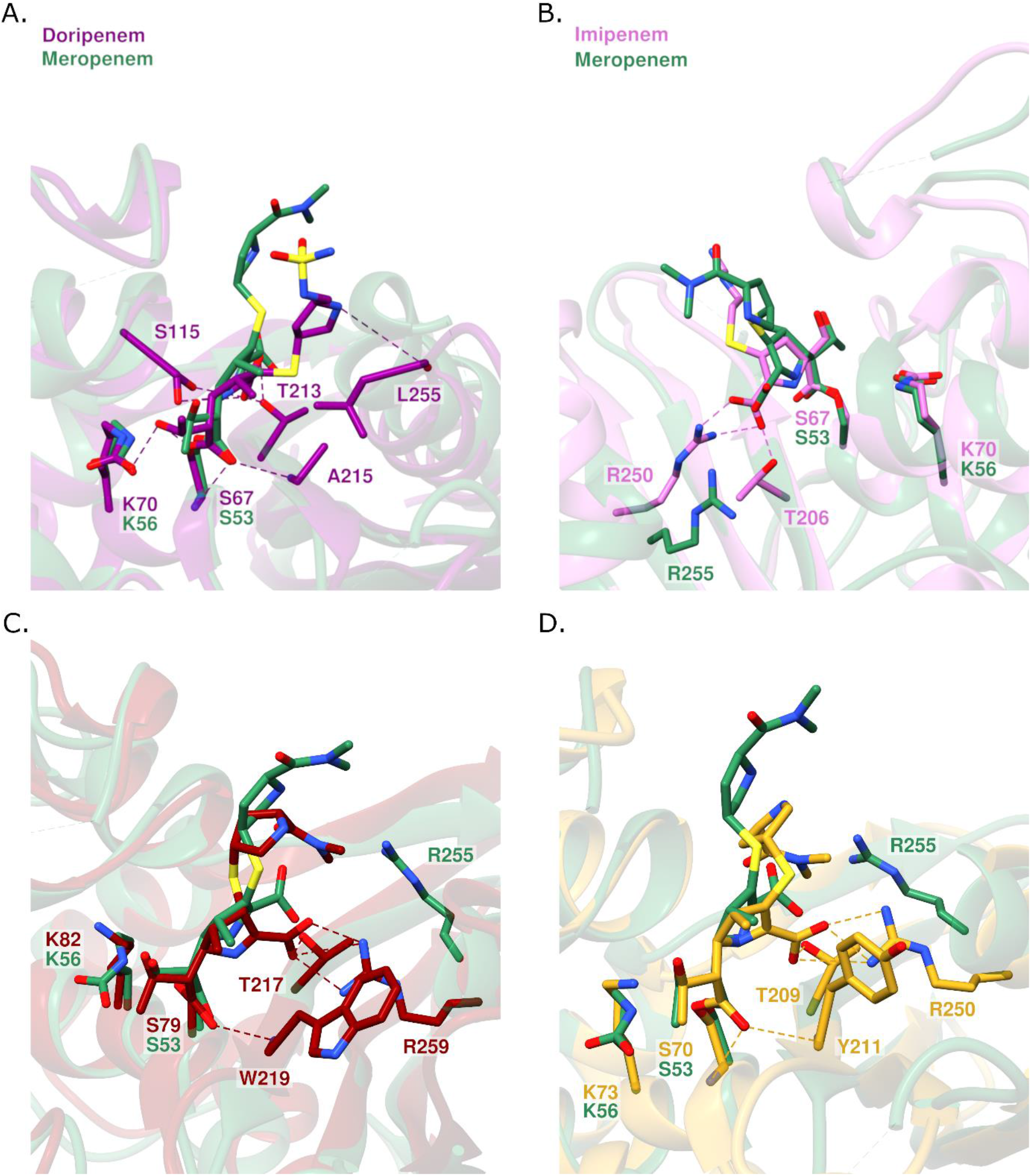
Carbapenem Binding to Class-D β-Lactamases. **a.** Overlay of OXA- 57:meropenem (green) and OXA-1:doripenem (PDB:3ISG^60^, purple) complexes. **b.** Overlay of OXA-57:meropenem (green) and OXA-10:imipenem (PDB:6SKP^67^, pink) complexes. **c.** Overlay of OXA-57:meropenem (green) and OXA-23:meropenem (PDB:4JF4^50^, red) complexes. **d.** Overlay of OXA-57:meropenem (green) and OXA-48:meropenem (PDB:6P98^49^, gold) complexes. For clarity, hydrogen bonds between OXA-57 and meropenem are not shown. Non-carbon atoms are colored by atom type and hydrogen bonding interactions are shown as dashed lines. RMSD values between structures are given in Supporting Information (**Table S9**). The figure was generated using UCSF Chimera^37^.

As is the case for the avibactam complexes described above, the most notable difference between the OXA-57:meropenem acyl-enzyme complex and comparator class D β- lactamase:carbapenem complexes is the observation that OXA-57 residue R255 is not favorably oriented to stabilize the meropenem acyl-enzyme. An arginine at the equivalent position is highly conserved in other CHDLs, and hydrogen bonds to the C3 carboxylate of bound carbapenems^48–50^. The OXA-10:imipenem complex contains two such bonds (**Figure 8b**) and the OXA-23:meropenem and OXA-48:meropenem complexes three (**Figure 8c**, **d**). Intriguingly OXA-1, which is not classed as a CHDL and is considered to be inhibited by carbapenems^60^, lacks an arginine at the equivalent position. However, this arginine residue does not appear essential for carbapenem hydrolysis, as the Gram-positive class-D β-lactamase BPU-1 also lacks arginine at this position but hydrolyzes carbapenems with comparable kinetic parameters to CHDLs from Gram-negative sources^61^ (**Table S2**).

Carbapenems are distinct from other β-lactams in the ability of their ring-opened species (acyl-enzymes or hydrolysis products) to access alternative tautomeric forms of their pyrroline ring: the Δ^2^ (enamine) and the Δ^1^ (imine, in both the (*S*)- Δ^1^ and (*R*)- Δ^1^ configurations)^18^. Previous studies with some ESBLs, such as SHV-1 and RTEM, have proposed the Δ^2^ enamine tautomer to be most relevant for efficient carbapenem hydrolysis^62,63^, and that efficient carbapenemases either minimize the extent of Δ^1^ tautomer formation or facilitate tautomerization of the Δ^1^-imine back to the Δ^2^-enamine^64^. In our OXA-57:meropenem structure, the meropenem acyl-enzyme was modelled as the Δ^2^ -enamine, based upon the location of the exocyclic sulfur. Various tautomeric states are crystallographically observed in the OXA:carbapenem complexes compared in **Figure 8**. In both the OXA-10:imipenem and OXA:48:meropenem complexes, bound carbapenems are modelled as the Δ^2^ -enamine tautomer. However, in the OXA-1:doripenem and OXA-23:meropenem structures, the carbapenem acyl-enzymes were modelled in the Δ^1^ -imine tautomer in the (*R*)- Δ^1^ and (*S*)- Δ^1^ configurations, respectively^65,66^. Considering these observations in the context of the reported kinetic parameters for carbapenem hydrolysis by each of these enzymes (**Table S2**), it is then clear that catalytic efficiency towards carbapenems does not necessarily correlate with the presence of a specific acyl-enzyme tautomer in the relevant crystal structures. In the case of OXA-57 our assignment of predominantly bound meropenem in the Δ^2^ -enamine tautomer, while consistent with the ability of the enzyme to hydrolyze meropenem, is not then an indicator of strong activity towards this substrate.

Access to the active site by the water molecule necessary for deacylation is fundamentally important to carbapenem hydrolysis by CHDLs ^15^. The carbamylated lysine is thought to act as a general base, activating the incoming water molecule, to attack the acyl-enzyme carbonyl carbon, ultimately liberating the inactivated product^14^. It has been proposed that steric hindrance during acylation, involving the carbapenem 6-α-hydroxyethyl group, causes expulsion of water from the active site necessitating subsequent recruitment of water from bulk solvent^16^. In the case of OXA-57, we found no evidence of solvent molecules positioned for deacylation in the active site of the uncomplexed enzyme, and there was no candidate deacylating water in the OXA-57:meropenem acyl-enzyme complex structure. While, given the comparatively low resolution of our crystallographic data, we cannot rule out the presence of a water molecule with insufficient occupancy to justify its inclusion in our final model, our MD simulations of the meropenem acyl-enzyme indicate only sporadic presence of an active site water molecule positioned for deacylation.

Different routes by which the deacylating water molecule can access the active site have been proposed for different CHDLs. In OXA-23, a hydrophobic patch formed by the conserved residues V128 and L166 shields the carbamylated lysine. Upon acylation, reorientation of the side chains of these residues causes formation of a channel close to the carbamylated lysine, allowing access of a water molecule^50,68^. In contrast, uncomplexed OXA-48 possesses a channel between the hydrophobic patch (residues V120 and L158) and the serine nucleophile that provides direct access of water to the carbamylated lysine, with acylation leading to only minimal conformational changes that widen this channel to promote water access^49^. In OXA- 57 V106 and I150 form an equivalent hydrophobic patch, but the OXA-57:meropenem acyl-enzyme complex structure provides no evidence of conformational changes leading to formation of a water access channel as described for OXA-23; and neither the uncomplexed or meropenem-bound OXA-57 structures contain a pre-existing solvent channel to the active site as reported for OXA-48.

We interrogated our MD simulations to investigate whether OXA-57 I150 might act as a “gating” residue regulating access of water to the active site. In 4 of our 5 repeat MD simulations I150 sampled different conformations throughout the trajectory (**Figure S19**), but there was no correlation between specific conformations and presence of water in the deacylating position, and the average RMSF of the I150 sidechain (1.0 Å) is consistent with the average RMSF values for all sidechains in the acyl-enzyme complex. Hence our simulations do not support a role for I150 facilitating or regulating access of water to the OXA- 57 active site to enable deacylation. Interestingly, the OXA-48 L158I mutant reduced catalytic efficiency towards meropenem, and also disfavored production of the alternative, β-lactone product as recently described for carbapenem hydrolysis by several Class D enzymes ^18,69^. This indicates that the presence of I150 in OXA-57 may be a contributing factor to its weak carbapenemase activity.

A noteworthy feature of some CHDLs, in particular enzymes such as OXA-23 and OXA-24 that originate in *Acinetobacter* spp., is formation of a hydrophobic bridge spanning the active site cleft. As first identified in OXA-24^51^ this involves contacts between the side chains of conserved residues flanking the active sight cleft, typically phenylalanine/tyrosine in the α3 - α4 loop and methionine/tryptophan on the β5 - β6 loop. In some cases, e.g. OXA-58, this bridge only forms upon ligand binding^52^. Candidate residues for involvement in hydrophobic bridge formation in OXA-57 are W86, on the α1 - α2 loop and Y206 on the β4 - β5 loop. However, the uncomplexed OXA-57 structure provides no evidence for a hydrophobic bridge between W86 and Y206, which lie 12 Å distant from one another, and conformational changes leading to formation of a hydrophobic bridge are not observed in the OXA-57:meropenem complex. Moreover, approach of W86 and Y206 within ≤ 4.0 Å of one another was rarely observed in MD simulations (**Figures S20, S21**), and these residues were generally further apart in simulations of the acyl-enzyme complex than in the uncomplexed enzyme. We conclude that, unlike equivalent complexes of some other CHDLs, OXA-57 cannot form a hydrophobic bridge across the active site cleft of the carbapenem acyl-enzyme complex.

The hydrophobic bridge plays a pivotal role in carbapenem hydrolysis by many CHDLs through constraining the tautomeric configuration of the carbapenem acyl-enzyme^70^. However, such a bridge is not present in all CHDLs, and carbapenem turnover occurs with comparable or even better efficiency in β-lactamases such as the OXA-48 group from which it is absent. In OXA-48, an alternative mechanism is proposed whereby a tyrosine (Y211) in an equivalent position to the methionine residue of the OXA-23 hydrophobic bridge orients towards the active site cleft forming a so-called “pseudobridge” that prevents formation of the (*R*)- Δ^1^ acyl-enzyme tautomer^49^. OXA-57 Y206 occupies the equivalent position to OXA-48 Y211, but is in a different orientation that is not consistent with a role in pseudobridge formation, and as described above makes only very transient interactions across the active site cleft during MD simulations. Thus, none of our findings support a role for OXA-57 Y206 in constraining carbapenem tautomerization, and our data do not identify any alternative mechanism by which this is achieved.

To investigate the broader implications of our observations, we carried out a genomic analysis of 1,295 *B. pseudomallei* clinical strains from Northeast Thailand^71^ and identified several chromosomal OXA isoforms, including OXA-59 along with other previously uncharacterized variants. These isoforms differ from OXA-57 by up to 4 single amino acid polymorphisms (SAPs). When the locations of these polymorphisms were mapped onto the OXA-57 crystal structure, most were found not in positions directly interacting with bound β-lactam substrates, and are thus unlikely to impact specificity or turnover (**Figure S22**). Indeed, previous kinetic studies identified minimal effects of specific substitutions, such as K231N in OXA-42 and D170N in OXA-59, on catalytic activity or substrate specificity^72^. However, two exceptions were identified: the S104P substitution in OXA-42 (not found in our strain collection) abolishes β-lactam hydrolysis *in vitro*, and the substitution of K56R in a variant type (VT8), observed in a single strain, could have functional implications. In summary, therefore, we expect the properties of OXA-57 described here to be representative of chromosomal OXA-β-lactamases from *B. pseudomallei*.

## 4. Conclusion

The data herein reveal *B. pseudomallei* OXA-57 to be a class-D β-lactamase capable of slow turnover of carbapenem as well as penicillin substrates, incapable of hydrolyzing (oxyimino) cephalosporins, and inhibited by clavulanate and (more weakly) avibactam. OXA-57 is monomeric (rather than dimeric as is the case for many other OXA enzymes) with an overall fold similar to those of other OXA enzymes, but with an extended, unstructured β4-β5 loop adjacent to the active site. Structures of complexes with avibactam and the acyl-enzyme of the carbapenem meropenem do not reveal substantial conformational changes associated with ligand binding, while a conserved active site arginine (OXA-57 R255) involved in ligand interactions of other OXA enzymes does not participate in hydrogen bonds to either the avibactam sulfate or meropenem C3 carboxylate groups. This is consistent with the relatively weak affinity of OXA-57 for both avibactam and meropenem. Molecular dynamics simulations of uncomplexed OXA-57 and its meropenem acyl-enzyme (assigned as the Δ^2^ tautomer) show reorientation of bound meropenem away from the position defined by the crystal structure, low occurrence of an active site water molecule positioned for deacylation and a lack of conformational changes, such as loop closure or hydrophobic bridge formation across the active site, associated with carbapenem hydrolysis by other class D β-lactamases. Thus, although kinetic data identify weak carbapenem-hydrolyzing activity by OXA-57, and the crystal structure indicates meropenem to be present in the (more hydrolytically labile) Δ^2^ - enamine tautomer, we do not identify specific features of OXA-57 that can be associated with carbapenemase activity. This is consistent with suggestions that weak carbapenemase activity is an inherent property of class D β-lactamases^53^. However, despite the weak *in vitro* activity of OXA-57, its coexistence with other resistance mechanisms, such as efflux pumps and alterations to permeability, and the potential for enhanced activity in the host cell through upregulation of expression and/or acquisition of point mutations (as has occurred in e.g. *Acinetobacter* sp.^73^), suggest that it should be considered as a potential contributing factor to β-lactam, particularly carbapenem, resistance in *B. pseudomallei*. Genomic analyses of chromosomal OXA genes in clinically derived strains of *B. pseudomallei* suggest our observations with OXA-57 are likely to be of general relevance. The unusual properties of OXA-57, as revealed by our combined biochemical and biophysical studies, also identify scope for improvement in β-lactamase inhibitors with potential application as part of new combination therapies for treating melioidosis.

## 5. Methods

### Full Methods are given in Supporting Information

#### Supporting information

Materials and Methods. Tables showing: PCR primers used for construct generation; comparison of carbapenem hydrolysis kinetics for Class-D β-lactamases, IC_50_ values for clavulanate, avibactam and vaborbactam inhibition of selected Class-D β-lactamases; X-ray data collection and refinement statistics; sequence and structural similarity of OXA-57 to other Class-D β-lactamases; B-factors for Lys56 carbamate; interactions of water molecule bridging meropenem and OXA-57 T202 in MD simulations; comparisons of selected OXA:avibactam and OXA:carbapenem complex structures. Figures showing proposed OXA-57 hydrolysis mechanism; purification of OXA-57; carbapenem hydrolysis by OXA-57; alternative conformations of carbamylated K56 in OXA-57:avibactam complex structure; starting models for molecular dynamics (MD) simulations; time series of backbone, core, K56 and meropenem RMSD values for MD simulations of uncomplexed OXA-57 and meropenem acyl-enzyme; backbone RMSF values for MD simulations of uncomplexed OXA-57 and OXA- 57:meropenem complex; Snapshots of structures of protein and bound meropenem during MD simulations of uncomplexed OXA-57 and OXA-57:meropenem; time series of hydrogen bond interactions made by meropenem during MD simulations of the OXA-57 complex; distribution of active site water molecules in MD simulations of OXA-57:meropenem complex; mobility of residue I150 during MD simulations of the OXA-57:meropenem complex, mobility of active site loops in MD simulations of uncomplexed OXA-57 and OXA-57:meropenem complex; sequence polymorphisms in *B. pseudomallei* chromosomal OXA β-lactamases.

#### Accession codes

Coordinates and structure factors for the OXA-57 crystal structures presented within the article have been deposited in the Protein Data Bank under accession codes 9HPT, 9HPU, 9HPW and 9HPY, for release on publication.

## Author Information Corresponding Author

James Spencer - School of Cellular and Molecular Medicine, University of Bristol, Bristol, BS8 1TD, UK

## Author contributions

É.C.B, C.K.C., C.J.S, N.C. and J.S. designed research; É.C.B, C.K.C, K.P., P.H, J.M.S, C.L.T and R.S performed research; É.C.B, C.K.C., K.P., P.H. and R.S analyzed data; N.C., A.J.M., C.J.S. and J.S. provided supervision; and É.C.B and J.S. drafted the manuscript with review and approval by all authors.

## Notes

The authors declare no conflicts of interest.

## Acknowledgements

Research was supported by the University of Bristol (studentship to É. C. B.) and the U.K. Biotechnology and Biological Sciences Research Council (grant no. BB/W001187/1 to J.S. and C.J.S.). This work is part of a project that has received funding from the European Research Council under the European Horizon 2020 research and innovation program (PREDACTED Advanced Grant Agreement no. 101021207) to A. J. M. and J. S. A.J.M. thanks the U.K. High-End Computing Consortium for Biomolecular Simulation, HECBioSim (http://hecbiosim.ac.uk), supported by EPSRC (grant no. EP/R029407/1). N.C. and J.S. acknowledge funding from the British Council through the Thai-UK World-class University Consortium. All simulations were conducted using the facilities of the Advanced Computing Research Centre at the University of Bristol (https://www.bris.ac.uk/acrc/). Diffraction data for uncomplexed OXA-57 (2.0Å resolution data set) were collected at the BL13-XALOC beamline at the ALBA Synchrotron with the collaboration of ALBA staff. We thank Diamond Light Source for beamtime (proposal no. MX23269) and the staff of beamline I04 for assistance with OXA-57: avibactam and OXA-57: meropenem data collections, and the staff of beamline I03 for assistance with uncomplexed OXA-57 (1.8 Å data set) data collection. We are also grateful to Stephen Hall for help with MATLAB scripting.

## Abbreviations

ESBL, extended-spectrum β-lactamases; IMAC, immobilized metal affinity chromatography; NH, no hydrolysis; BLI, β-lactamase inhibitor; DBO, diazabicyclooctane; IC_50_, half maximal inhibitory concentration; RMSD, root-mean-squared deviation; RSCC, real-space correlation coefficient; RMSF, root-mean square fluctuation; CHDL, carbapenem-hydrolyzing class-D β- lactamases; MIC, minimum inhibitory concentration.

## Data Availability

Starting models and representative snapshot structures from MD simulations will be made freely available at the University of Bristol Research Data Repository (https://data.bris.ac.uk/).

## Notes

### Competing Interest Statement

The authors have declared no competing interest.

