## Supplementary Information for "Structure and dynamics of *Burkholderia pseudomallei* OXA-57, a distinctive low efficiency class D β-lactamase with carbapenem-hydrolyzing activity"

Supporting information

*Éilís C. Bragginton [1,2†], Charlotte K. Colenso [1,3], Karina Calvopiña [4], Philip Hinchliffe [1], John M. Shaw [1], Catherine. L. Tooke [1,5†], Rathanin Seng [6,7], Narisara Chantratita [6,8], Adrian J. Mulholland [3], Christopher J. Schofield [4] and James Spencer [1*]*

[1] School of Cellular and Molecular Medicine, University of Bristol, Bristol, BS8 1TD, U.K.

[2] Diamond Light Source, Harwell Science and Innovation Campus, Didcot OX11 0DE, U.K.

[3] Centre for Computational Chemistry, School of Chemistry, University of Bristol, Bristol, BS8 1TS, U.K.

[4] Chemistry Research Laboratory, Department of Chemistry and the Ineos Oxford Institute for Antimicrobial Research, University of Oxford, Oxford, OX1 3TA, U.K.

[5] Department of Life Sciences, University of Bath, Bath, BBA2 7AY, U.K.

[6] Department of Microbiology and Immunology, Faculty of Tropical Medicine, Mahidol University, Bangkok, Thailand.

[7] Department of Medical Science, Amnat Charoen Campus, Mahidol University, Amnat Charoen, Thailand.

[8] Mahidol Oxford Tropical Medicine Research Unit, Faculty of Tropical Medicine, Mahidol University, Bangkok, Thailand.

[†]Present address

**Supplementary Methods**

Reagents were purchased from Merck unless otherwise specified.

### OXA-57 purification

DNA encoding for the predicted mature polypeptide of OXA-57 (codons 24 to 269) was cloned into pET28b (Novagen) (**Table S1**) and expressed in *Escherichia coli* BL21 (DE3) cells (Novagen). Cells carrying this plasmid were grown in LB broth, supplemented with 50 µg $ml^{-1}$ kanamycin, at 37 ºC to an optical density of 0.6 at 600 nm ($OD_{600}$). Cells were subsequently grown at 18 ºC overnight with the addition of 1 mM isopropyl β-D-1-thiogalactopyranoside (IPTG) to induce expression. Cells were harvested by centrifugation (2 700 x*g*, 15 min) and resuspended in 50 mM Tris pH 8.0, 20 mM Imidazole pH 8.0, 500 mM NaCl with complete EDTA-free protease inhibitor (Roche). Cells were lysed by 3 passages through a French press cell disruptor (900 psi) and cell debris removed by centrifugation (50 000 x*g*, 1 hr, 4 ºC). The supernatant was passed through a 0.45 µm filter prior to loading onto a pre-equilibrated 5 ml HisTrap column (Cytiva Life Sciences). The column was equilibrated and washed with 10 column volumes of 50 mM Tris pH 8.0, 20 mM Imidazole pH 8.0, 50 mM NaCl and eluted with a linear imidazole gradient (20 – 300 mM) over 36 column volumes. The purity of eluted fractions was assessed by SDS-PAGE. Fractions containing OXA-57 were pooled and buffer exchanged into 20 mM Tris pH 8.4, 150 mM NaCl, 2.5mM $CaCl_2$ in an Amicon 10-kDa molecular weight cutoff (MWCO) centrifugal filter. The N-terminal His-tag was removed from OXA-57 with the addition of thrombin (15 µl, 50x units) (Novagen) and incubated overnight at room temperature. Cleaved OXA-57 was loaded onto a HiLoad 16/600 Superdex 75 pg column (Cytiva Life Sciences), pre-equilibrated in 50 mM HEPES pH 7.5, 150 mM NaCl, and peak fractions assessed by SDS-PAGE. Fractions with >90 % purity were pooled and concentrated to 15 mg $ml^{-1}$ using an Amicon 10-kDa molecular weight cutoff (MWCO) centrifugal filter. The concentration of OXA-57 was determined by measuring absorbance at $\lambda_{280\ nm}$ using the protein extinction coefficient ($\varepsilon$ = 56 045 $M^{-1}$ $cm^{-1}$) calculated using the ExPASy ProtParam tool. Purified OXA-57 was stored in 50 mM HEPES pH 7.5, 150 mM NaCl supplemented with 20 % glycerol at -70 ºC.

**Enzyme assays**

Enzyme assays for all substrates except FC5 were followed at 25 ºC in 100 mM sodium phosphate pH 7.0, 150 mM $NaCHO_3$ (Buffer A) with 1 µM OXA-57. Steady-state kinetic parameters were determined with an Tecan Infinite M200 pro plate reader (Tecan) using Greiner half area 96-well plates. Hydrolysis rates were measured using the following extinction coefficients ($\Delta\varepsilon$) and concentration ranges: ampicillin $\Delta\varepsilon$, -809 $M^{-1}$ $cm^{-1}$ at 235 nm (37.5 – 600 µM), aztreonam $\Delta\varepsilon$, -700 $M^{-1}$ $cm^{-1}$ at 320 nm (37.5 – 600 µM), cephalothin $\Delta\varepsilon$, -8790 $M^{-1}$ $cm^{-1}$ at 265 nm (9.4 – 300 µM), cefotaxime $\Delta\varepsilon$, -7250 $M^{-1}$ $cm^{-1}$ at 265 nm (18.8 – 300 µM), ceftazidime $\Delta\varepsilon$, -7455 $M^{-1}$ $cm^{-1}$ at 265 nm (18.8 – 300 µM), imipenem $\Delta\varepsilon$, -10930 $M^{-1}$ $cm^{-1}$ at 298 nm (9.4 – 300 µM), meropenem $\Delta\varepsilon$, -7200 $M^{-1}$ $cm^{-1}$ at 298 nm (9.4 – 300 µM). Steady-state velocities were calculated from the initial rates of substrate hydrolysis and plotted against substrate concentration. Kinetic parameters were calculated and analyzed using GraphPad Prism 9.0. Nonlinear regression of the data determined the $K_M$ and $k_{cat}$ as described by the Michaelis-Menten equation:

$$v = \frac{V_{max}[S]}{K_M + [S]} \quad (1)$$

Where $V_{max}$ can be described as a function of $k_{cat}$ multiplied by the enzyme concentration [E]:

$$V_{max} = k_{cat} \times [E] \quad (2)$$

For some substrates, inhibition was observed at high substrate concentrations. In these cases, a modified Michaelis-Menten equation was used to determine the kinetic parameters considering substrate inhibition:

$$v = \frac{V_{max}[S]}{(K_M + [S]) \times (1 + \frac{[S]}{k_i})}$$

(3)

Steady-state kinetic parameters for the fluorogenic substrate FC5 were determined using 12.5 nM OXA-57 in 100 mM phosphate buffer, pH 7.4, 0.01% v/v Triton X-100, 150 mM $NaHCO_3$ (Buffer B) over a concentration range of 0.009 – 20 µM. The assays were conducted at room temperature in clear-bottomed Greiner 384 black well microplates and fluorescence intensity was monitored at λex = 380 nm and λem = 460 nm in a ClarioStar or PHERAstar FS microplate reader (BMG LabTech). Kinetic parameters were calculated as described above.

The $IC_{50}$'s of inhibitors were determined using a well-described fluorogenic assay by monitoring the enzymatic breakdown of FC5[1]. In brief, inhibitor compounds at concentrations ranging from $2x10^{-4}$ – 4 mM were incubated with OXA-57 (12.5 nM) in buffer B for 10, 30 and 60 min, followed by the addition of FC5 (5 µM). The assays were conducted as described above for the determination of FC5 kinetic parameters. Normalized data were fitted using a four-parameter function: log (inhibitor) vs. response, variable slope in GraphPad Prism 6 to obtain $IC_{50}$ values.

**Crystallization and structure refinement**

OXA-57 was crystallized using the vapor diffusion method. Sitting drops were prepared in a 1:1 ratio of crystallization reagent to 10 μg $ml^{-1}$ OXA-57 in 50 mM HEPES pH 7.5, 150 mM NaCl.

The 2.0 Å uncomplexed OXA-57 structure, OXA-57:meropenem and OXA-57:avibactam complex structures (PDB: 9HPT, 9HPW and 9HPY) were crystallized in 0.12 M Monosaccharides (0.2 M D-glucose, 0.2 M D-mannose, 0.2 M D-galactose, 0.2 M L-fucose, 0.2 M D-xylose, 0.2 M N-acetyl-D-glucosamine), 0.1 M Tris; BICINE pH 8.5, 37.5 % (v/v) Precipitant mix (25 % (v/v) MPD, 25 % (w/v) PEG 1000, 35 % (w/v) PEG 3350) (from the Morpheus crystallization screen, Molecular Dimensions). Drops (2 µl protein solution + 2 µl reservoir solution) were equilibrated against 500 µl reservoir solution in CrysChem 24-well plates (Hampton Research) and incubated at 18 ºC. The 1.8 Å uncomplexed OXA-57 structure (PDB: 9HPU) was crystallized in 0.2 M ammonium acetate, 0.1 M Bis-Tris: HCl pH 6.5, 25% (w/v) PEG 3350 (TOP96, Anatrace). Drops (0.2 µl protein solution + 0.2 µl reservoir solution) were prepared in MRC 2 Lens crystallisation plates (SWISSCI) and incubated as above.

For inhibitor and substrate soaking, the 2.0 Å uncomplexed OXA-57 crystals were soaked in mother liquor supplemented with 2 mM Avibactam or 10 mM meropenem. All crystals were exposed to mother liquor and flash-frozen in liquid nitrogen prior to data collection. Diffraction data for the 2.0 Å uncomplexed OXA-57 crystals were collected at the Alba synchrotron on beamline BL13 – XALOC. Images were indexed and integrated with XDS and scaled using Aimless (CCP4 suite[2]). For the 1.8 Å uncomplexed, inhibitor- and substrate-soaked crystals, diffraction data was collected at Diamond Light Source on beamlines i04 and i03 respectively. Images were indexed and integrated using the Dials[3] and Xia2[4] pipeline at Diamond Light Source and subsequently scaled in Aimless (CCP4 suite[2]). All data were phased by molecular replacement using Phaser[5] (CCP4 suite[2]). For the 2.0 Å uncomplexed OXA-57 structure, the PDB identifier 3W40 (YdxI from *Burkholderia thailandensis*) was used. For the 1.8 Å uncomplexed, inhibitor- and substrate-soaked data, the 2.0 Å uncomplexed OXA-57 structure was used as the starting model for molecular replacement. The resulting models were iteratively refined using phenix.refine[6] and manually model building in WinCoot[7]. Geometry restraints for Avibactam and Meropenem were calculated using eLBOW and omit maps generated in phenix[6] from the final model in the absence of ligand.

**Molecular modelling and simulations**

Unstructured regions of the uncomplexed OXA-57 (PDB entry 9HPT, resolution 2.05 Å) and OXA-57:meropenem complex (PDB entry 9HPW resolution 2.42 Å) X-ray crystal structures (loops β4-β5 and β6-α8, and α1-α2, β4-β5 and β6-α8, respectively), were built using MODELLER v9.24[8]. In the case of the β4-β5 loop, the X-ray crystal structure of the class D β-lactamase Ybxl (*Burkholderia thailandensis)* was used as a template since this region is defined (PDB entry 6NI0, resolution 2.30 Å; 100% sequence identity between OXA-57 residues L209-R214). PROCHECK was used to check the quality of the resulting structures[9]. Protonation states of titratable groups were predicted using the H++ server v3.2 (http://biophysics.cs.vt.edu/H++)[10] at pH7.5. For the acyl-enzyme complex, partial atomic charges for the $\Delta^2$ tautomer of meropenem were generated using restrained electrostatic potential (RESP) fitting as implemented in the R.E.D. Server Development 2.0[11] following geometry optimization at the HF/6-31G* level using Gaussian 16. Missing forcefield parameters were taken from analogous GAFF parameters. The carbamylated lysine (K56) in uncomplexed and acyl-enzyme structures was parameterized with the same protocol. In both systems, all crystallographic waters were retained, and a disulfide bond was set by linking C27 and C45.

All simulations were prepared using the LEaP module in AmberTools18[12]. Proteins were solvated in a truncated octahedron with TIP3P water molecules having a 10 Å minimum distance between any protein atom and the octahedron edge. Chloride ions were added to attain a neutral net charge along with an additional 150 mM NaCl concentration. Systems were minimized then heated to 300 K at constant volume over 200 ps. Short equilibration dynamics totaling 2 ns were run at constant pressure with isotropic position scaling at 1 bar, controlled by the Berensden barostat with a pressure relaxation time of 1 ps. The temperature was maintained using Langevin dynamics with the collision frequency set to 5 $ps^{-1}$. Protein backbone atoms (N, Cα, C) and heavy atoms of K56 and meropenem were constrained by a harmonic potential with a force constant of 10 kcal/mol – $Å^2$ which was gradually reduced over the minimization, heating, and equilibration steps. The Particle Mesh Ewald (PME) method was used to calculate long-range electrostatic interactions while the short-range electrostatic and van der Waals interactions were truncated at 10 Å. All bond lengths involving hydrogen were constrained using the SHAKE algorithm. Unrestrained molecular dynamics simulations were performed for 500 ns with a timestep of 2 fs and coordinates were saved every 10 ps. All simulations were repeated five-fold (R1-R5) and run on the University of Bristol supercomputer BlueCrystal 4. The pmemd.cuda module of AMBER18 was used to run the simulations using the ff14SB forcefield[13] at 300 K under periodic boundary conditions.

All simulations were analyzed using the *cpptraj* module within AmberTools18 having removed the first 75 ns of each trajectory to account for equilibration, unless otherwise stated. Mass-weighted root-mean-squared deviation (RMSD) calculations were computed with reference to the crystal structure. Average structures were calculated over 425 ns using N, Cα, and C atoms, and used as reference in root-mean-squared fluctuation (RMSF) calculations. Representative structures in Figures 4 -6 are those having the lowest RMSD to the average structure calculated over all repeat simulations. For hydrogen bonding analyses the acceptor to donor heavy atom distance cut-off was set to 3.5 Å. Trajectories were visualized and manipulated using VMD[14]. UCSF Chimera[15], gnuplot and MATLAB were used to produce figures. The *mindist* function of AMBER20 was used to generate Supplementary Figures S20 and 21.

**Table S1.** Primers used to generate recombinant vector pET28b (+) *OXA-57* (Δ1-69). Primers were supplied by Eurofins genomics.

| Primers | Sequence |
|---|---|
| Forward | CGCGCGGCAGCCATATGAAGACTATCTGCACGGCTATTGCG |
| Reverse | GCTCGAATTCGGATCCTTAGCGGGCTGCGAGTAAGCG |

**Table S2.** Comparison of Class-D β-lactamase steady-state kinetics for imipenem (IMI) and meropenem (MER) hydrolysis.[16-18]

| Class-D β-lactamase | Organism | $K_M$ (μM) | | $k_{cat}$ ($s^{-1}$) | | $k_{cat}/K_M$ ($M^{-1}$ $s^{-1}$) | |
|---|---|---|---|---|---|---|---|
| | | IMI | MER | IMI | MER | IMI | MER |
| OXA-57 | *B. pseudomallei* | 51.0 | 21.0 | 0.04 | 0.01 | $7.8 \times 10^2$ | $3.3 \times 10^3$ |
| OXA-2[1] | *P. aeruginosa* | ≤2.0 | ≤2.0 | 0.18 | 0.11 | $\geq 9.0 \times 10^4$ | $\geq 5.5 \times 10^4$ |
| OXA-10[1] | *P. aeruginosa* | 2.0 | 5.6 | 0.039 | 0.041 | $2.1 \times 10^4$ | $7.0 \times 10^3$ |
| OXA-23[2] | *A. baumanii* | 4.8 | ≤1.0 | 0.35 | 0.068 | $7.4 \times 10^5$ | $\geq 6.8 \times 10^4$ |
| OXA-24[1] | *A. baumanii* | ≤2.0 | ≤2.0 | 1.7 | 0.11 | $\geq 8.5 \times 10^5$ | $\geq 5.5 \times 10^4$ |
| OXA-48[1] | *K. pneumoniae* | 5.3 | ≤2.0 | 4.8 | 0.16 | $1.3 \times 10^6$ | $\geq 8.0 \times 10^4$ |
| OXA-58[1] | *A. baumanii* | 6.0 | ≤2.0 | 1.8 | 0.019 | $2.9 \times 10^5$ | $\geq 9.5 \times 10^3$ |
| BPU-1[3] | *B. pumilus* | 6.6 | 4.2 | 0.26 | 0.014 | $6.2 \times 10^3$ | $3.3 \times 10^3$ |

**Table S3**. Comparison of $IC_{50}$ values for Clavulanic acid, Avibactam and Vaborbactam inhibition of selected Class-D β-lactamases[19-23].

| Class-D β-lactamase | Organism | Clavulanate | | Avibactam | | Vaborbactam | |
|---|---|---|---|---|---|---|---|
| | | Incubation time (min) | $IC_{50}$ (μM) | Incubation time (min) | $IC_{50}$ (μM) | Incubation time (min) | $IC_{50}$ (μM) |
| OXA-57 | *B. pseudomallei* | 10<br>30 | 30.3<br>12.3 | 10<br>30 | 58.8<br>18.8 | - | |
| OXA-10 | *P. aeruginosa* | NA | *< 40[4] | 10<br>30 | 0.17[6]<br>0.04[6] | NA | > 400[7] |
| OXA-23 | *A. baumanii* | - | | 10<br>30 | 0.09[6]<br>0.17[6] | 10 | * 120[8] |
| OXA-48 | *K. pneumoniae* | 30 | *16[5] | 10<br>30 | 0.50[6]<br>0.14[6] | NA | 32[7] |

* Reaction not supplemented with $NaHCO_3$

NA: not available.

**Table S4.** X-ray diffraction data collection and refinement statistics.

| | Uncomplexed OXA-57<br><br>PDB: 9HPT | Uncomplexed OXA-57<br><br>PDB: 9HPU | OXA-57: avibactam<br><br>PDB: 9HPY | OXA-57: meropenem<br><br>PDB: 9HPW |
|---|---|---|---|---|
| **Data Collection** | | | | |
| Space Group | C 2 2 21 | C 2 2 21 | C 2 2 21 | C 2 2 21 |
| **Cell dimensions** | | | | |
| *a, b, c* (Å) | 66.22, 111.56, 66.20 | 65.79, 110.35, 64.80 | 65.78, 110.84, 66.16 | 65.93, 109.86, 66.36 |
| α, β, γ (º) | 90, 90 ,90 | 90, 90, 90 | 90, 90 ,90 | 90, 90 ,90 |
| Wavelength (Å) | 0.9 | 0.98 | 0.91 | 0.91 |
| Resolution (Å) | 43.17 – 2.0 (2.05 – 2.0) | 42.59 – 1.80 (1.84 – 1.80) | 55.42 – 2.48 (2.58 – 2.48) | 56.53 – 2.42 (2.51 – 2.42) |
| $R_{pim}$ | 0.020 (1.041) | 0.049 (0.917) | 0.048 (0.732) | 0.036 (0.568) |
| CC1/2 | 1.0 (0.532) | 0.998 (0.38) | 0.999 (0.59) | 0.997 (0.667) |
| $I/\sigma$ ($I$) | 17.3 (0.7) | 8.9 (0.8) | 9.0 (0.8) | 11.7 (1.2) |
| Completeness (%) | 100.0 (100.0) | 100.0 (100.0) | 100.0 (100.0) | 99.7 (99.3) |
| Redundancy | 13.1 (12.0) | 11.4 (12.7) | 12.3 (12.9) | 12.8 (12.8) |
| **Refinement**<br>Resolution (Å) | 43.17 – 2.0 | 42.01 – 1.80 | 55.42 – 2.48 | 56.53 – 2.42 |
| No. Reflections | 16959 | 22237 | 8825 | 9432 |
| $R_{work}/R_{free}$ | 0.2072/0.2311 | 0.1890/0.2143 | 0.1925/0.2451 | 0.1971/0.2601 |
| **No. non-H atoms** | | | | |
| Protein | 1831 | 1810 | 1812 | 1800 |
| Solvent | 24 | 134 | 9 | 12 |
| Ligand | - | 1 | 17 | 26 |
| ***B*-factors (Å$^2$)** | | | | |
| Protein | 65.6 | 36.2 | 76.4 | 79.9 |
| Solvent | 53.8 | 39.6 | 74.3 | 71.2 |
| Ligand | - | 42.0 | 71.1 | 114.6 |
| **r.m.s deviations** | | | | |
| Bond lengths (Å) | 0.007 | 0.006 | 0.008 | 0.007 |
| Bond angles (º) | 0.832 | 0.830 | 1.01 | 0.88 |
| **Ramachandran (%)** | | | | |
| Outliers | 0.44 | 0.00 | 0.88 | 1.35 |
| Favored | 94.27 | 96.43 | 91.59 | 93.27 |

**Table S5.** Similarity of OXA-57 to other class-D β-lactamases.

| Protein | PDB accession code | Resolution (Å) | Sequence identity (%) | Cα RMSD (Å) |
|---|---|---|---|---|
| OXA-1 | 1M6K | 1.50 | 34.67 | 1.41 |
| OXA-10 | 1FOF | 2.00 | 25.36 | 1.76 |
| OXA-23 | 4K0X | 1.61 | 22.66 | 1.50 |
| OXA-24 | 3G4P | 1.97 | 18.75 | 1.62 |
| OXA-48 | 4S2P | 1.70 | 27.36 | 1.62 |
| OXA-58 | 4OH0 | 1.30 | 21.95 | 1.81 |

**Table S6.** B-factors for OXA-57 Lys56 carbamate.*

| | Uncomplexed OXA-57<br><br>PDB: 9HPT | Uncomplexed OXA-57<br><br>PDB: 9HPU | OXA-57: avibactam<br><br>PDB: 9HPY | OXA-57: meropenem<br><br>PDB: 9HPW |
|---|---|---|---|---|
| Atom | B-factor (Å$^2$) | B-factor (Å$^2$) | B-factor (Å$^2$) | B-factor (Å$^2$) |
| KCX56 Nε | 50.30 | 27.75 | 66.83 | 76.69 |
| KCX56 CX | 54.01 | 28.90 | 67.38 | 76.41 |
| KCX56 OQ1 | 51.26 | 27.57 | 69.48 | 76.94 |
| KCX56 OQ2 | 50.47 | 27.36 | 74.43 | 76.68 |

*Lysine-arbamate occupancy was set to 1.0 in all cases.

**Table S7.** Bridging water interactions between the C3 carboxylate group of meropenem and T202. Values correspond to the number of frames (total 42,501) a bridging water is present.

| Water-mediated acceptor:donor/acceptor | R1 | R2 | R3 | R4 | R5 |
|---|---|---|---|---|---|
| MEM53@*O01*:T202@*OG1* | 5787 | | | 4384 | |
| MEM53@*O01*:T202@*HG1* | 5197 | 12136 | 7952 | 6649 | 13194 |
| MEM53@*O03*:T202@*OG1* | 10709 | | 5417 | | |
| MEM53@*O03*:T202@*HG1* | 6359 | 6428 | 7159 | 6255 | 6033 |
| MEM53@*O01*:MEM53@*H01*:T202@*OG1* | | 12062 | 10039 | | 10026 |
| MEM53@*O03*:MEM53@*H01*:T202@*OG1* | 6094 | 8446 | 12298 | | 5412 |

**Table S8.** Similarity of OXA-57:avibactam structure to other OXA:avibactam complexe structures.

| Protein | PDB accession code | Resolution (Å) | Cα RMSD (Å) |
|---|---|---|---|
| OXA-10:avibactam | 4S2O | 1.70 | 1.72 |
| OXA-24:avibactam | 4WM9 | 2.40 | 1.56 |
| OXA-48:avibactam | 4S2N | 2.00 | 1.61 |

**Table S9.** Similarity of OXA-57:meropenem structure to other OXA:carbapenem complexes.

| Protein | PDB accession code | Resolution (Å) | Cα RMSD (Å) |
|---|---|---|---|
| OXA-1:doripenem | 3ISG | 1.40 | 1.45 |
| OXA-10:imipenem | 6SKP | 1.89 | 1.70 |
| OXA-23:meropenem | 4JF4 | 2.14 | 1.58 |
| OXA-48:meropenem | 6P98 | 1.75 | 1.58 |

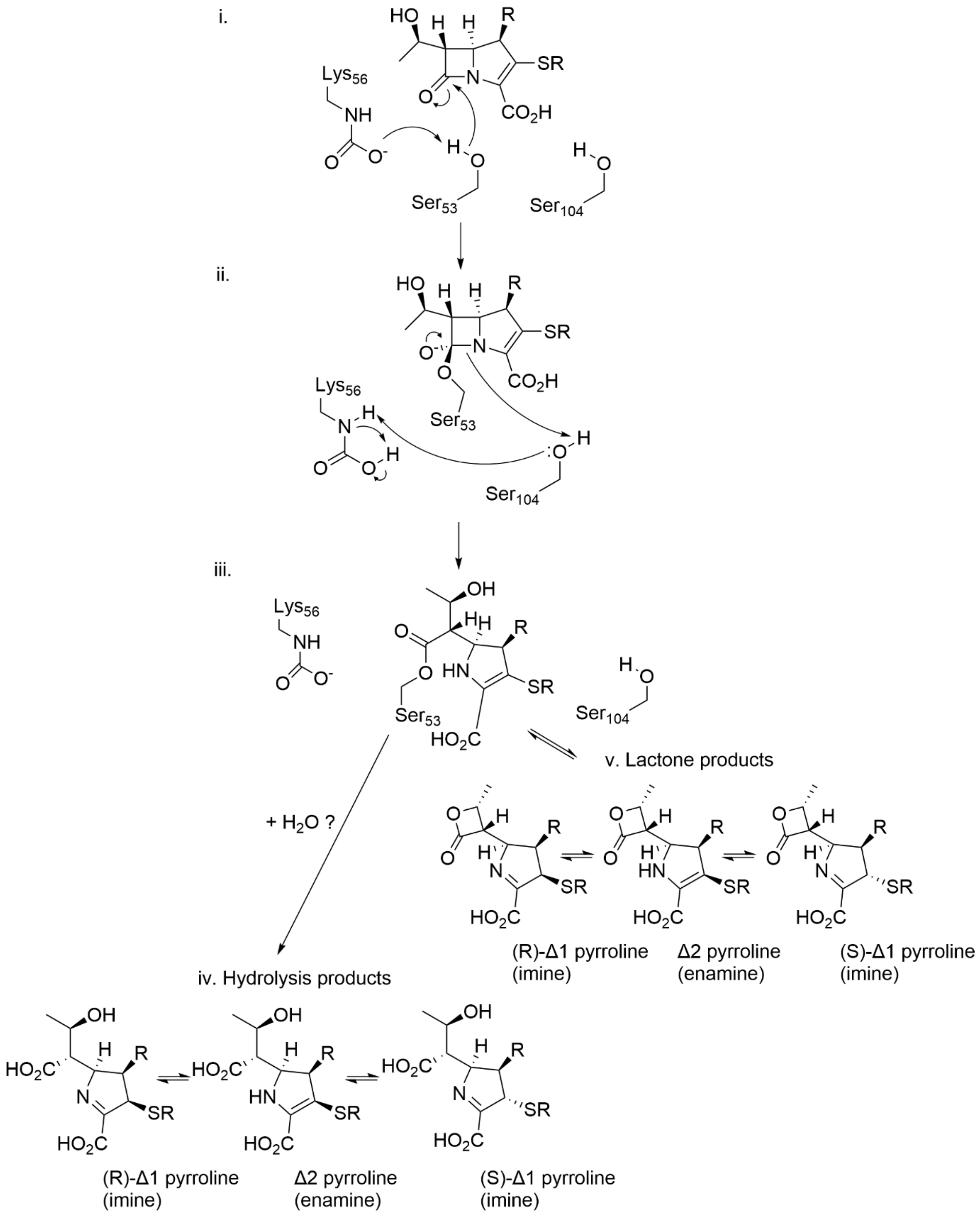


**Figure S1. Possible mechanisms of reaction of carbapenem substrates with OXA β-lactamases. i.** Ser53 is activated for nucleophilic attack on the β-lactam carbonyl by the carbamylated Lys56. **ii.** Nucleophilic attack results in a tetrahedral acylation intermediate that resolves to the ring-opened acylenzyme **iii.** *via* protonation of the amide nitrogen by Ser104. The acylenzyme can resolve by hydrolysis **iv**. through the action of incoming deacylating water molecule to yield products in alternative tautomeric and epimeric forms. The alternative lactone products **v.** are reversibly generated through acylenzyme rearrangement and, like the hydrolyzed products, can also adopt different tautomeric and epimeric states.

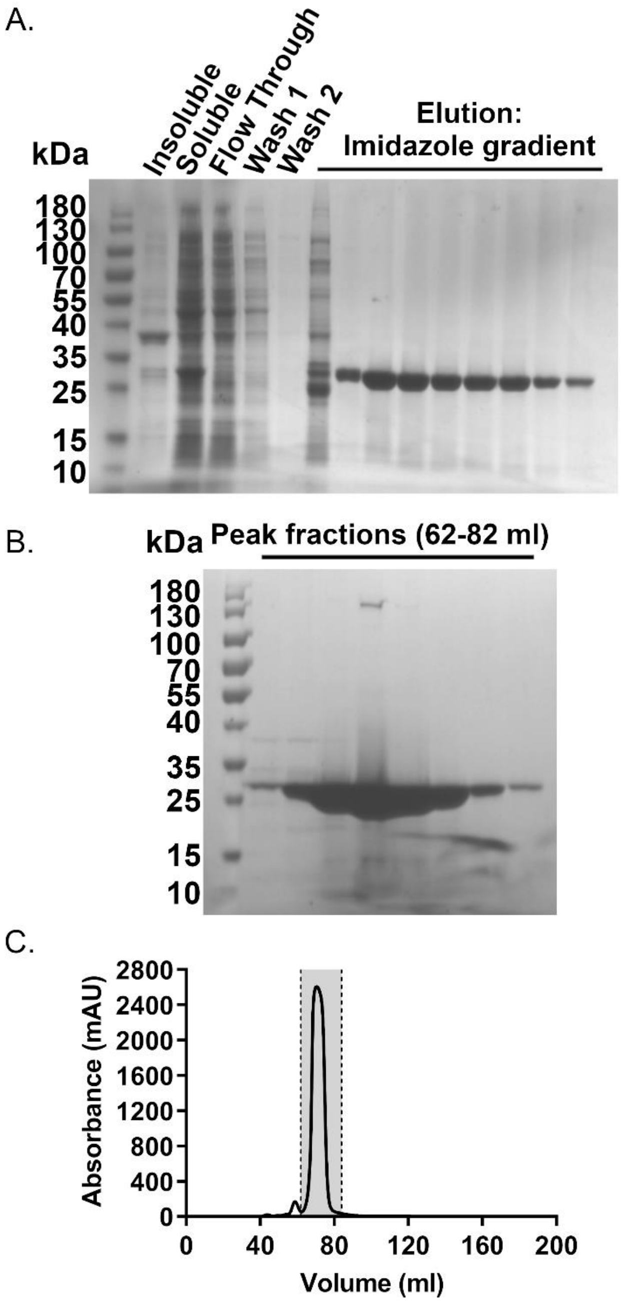


**Figure S2. Purification of recombinant OXA-57.** A. SDS-PAGE showing fractions from IMAC purification with elution on a linear imidazole gradient. B. SDS-PAGE showing peak fractions from Size Exclusion Chromatography (SEC) on Superdex 75 column. C. Representative chromatogram of HiLoad 16/600 Superdex 75 pg SEC monitored by absorbance at 280 nm. Shaded gray region of the chromatogram represents the peak fractions shown in B, OXA-57 elutes as a single symmetrical peak of approximately 27 kDa molecular mass.

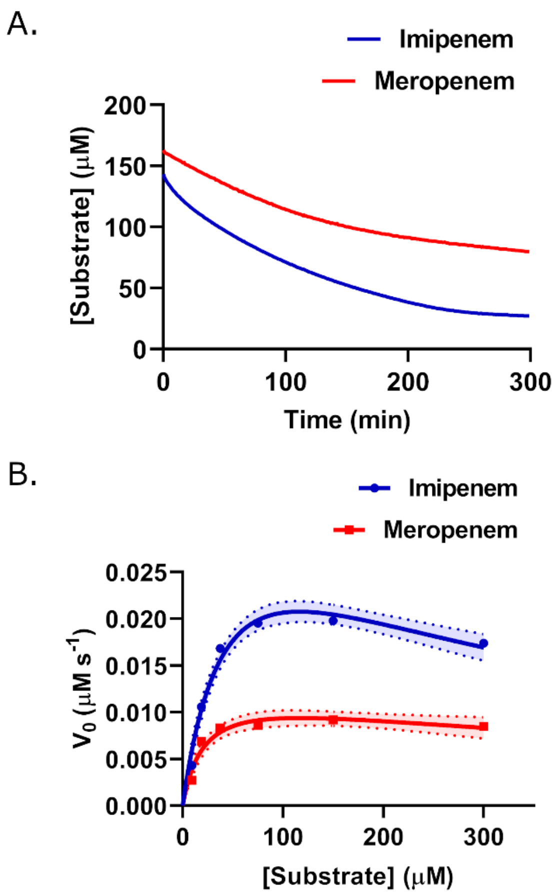


**Figure S3. Hydrolytic activity of OXA-57 against carbapenems.** A. Representative progress curves of imipenem (blue) and meropenem (red) hydrolysis at 150 μM with 1 μM OXA-57. B. Michaelis-Menten plots of imipenem (blue) and meropenem (red) hydrolysis kinetics. Each measurement was carried out in triplicate, dotted lines represent the 95 % confidence bands from nonlinear regression fitting.

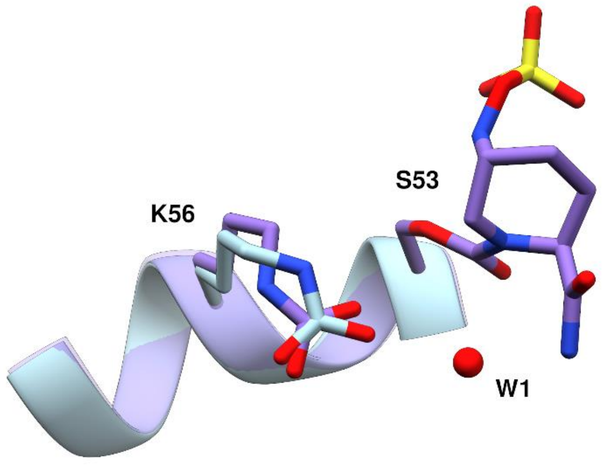


**Figure S4. Change in orientation of carbamylated lysine K56 upon avibactam binding.** OXA-57 protein chain (cartoon) and K56 side chain (sticks) are shown in light blue and purple in the uncomplexed and OXA-57:avibactam structures, respectively. Bound avibactam is shown as sticks, with carbon atoms purple and non-carbon atoms colored by atom type. Water molecules shown as red spheres.

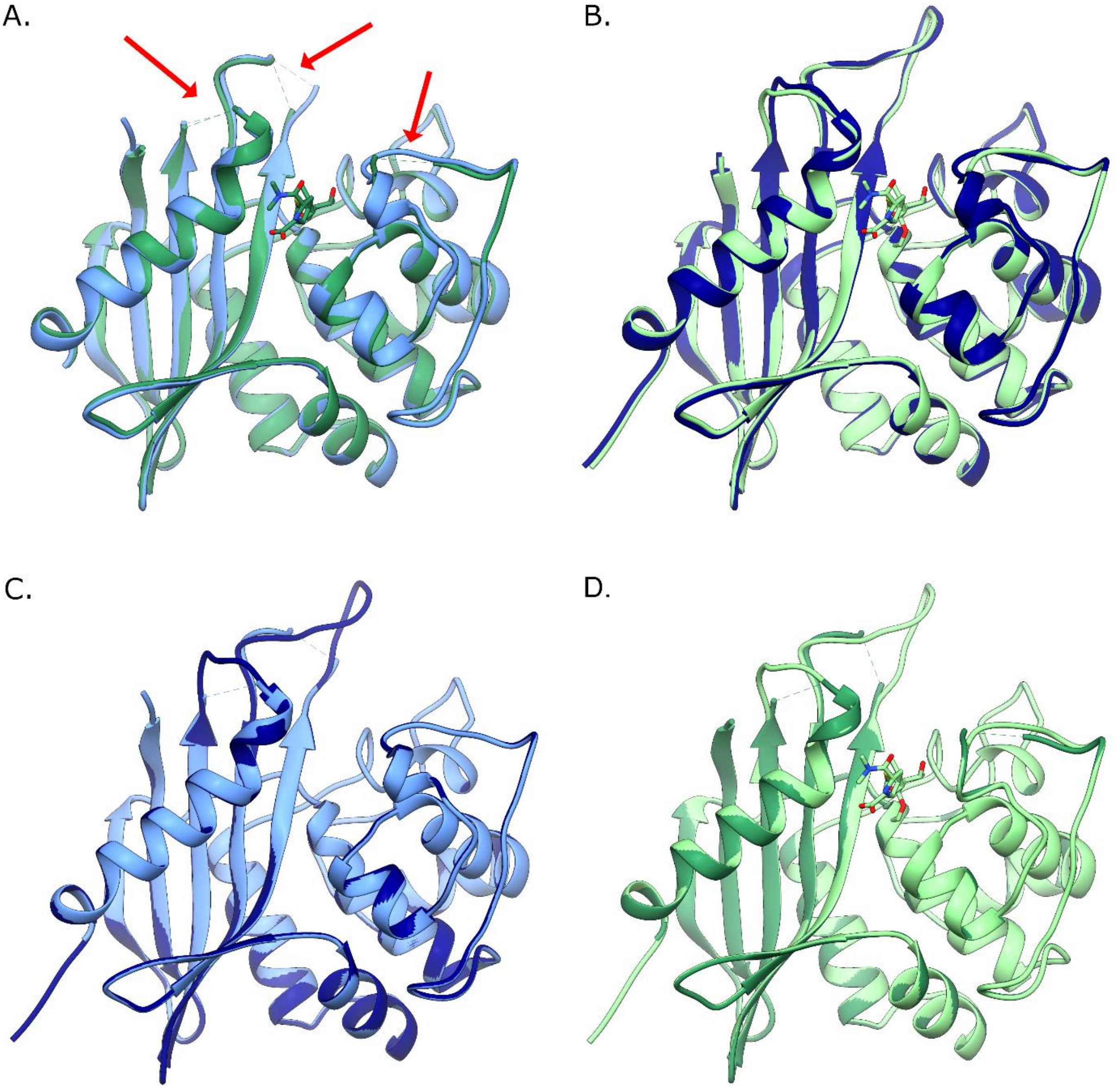


**Figure S5. OXA-57 Structures Used for Molecular Dynamics Simulations.** A. X-ray crystal structures (this work) of uncomplexed OXA-57 (PDB: 9HPT; blue) and OXA-57: meropenem complex (PDB: 9HPW; green). Red arrows indicate positions where unmodelled loops in the crystal structures were built for simulations. B. Models of uncomplexed OXA-57 (blue) and meropenem complex (mint) with β4 - β5 and β6 - α8, and α1 - α2, β4 - β5 and β6 - α8 loops, respectively, built using MODELLER. C. Uncomplexed OXA-57 crystal structure (blue) and after addition of modelled loops (navy). D. OXA-57:meropenem complex crystal structure (green) and after addition of modelled loops (mint).

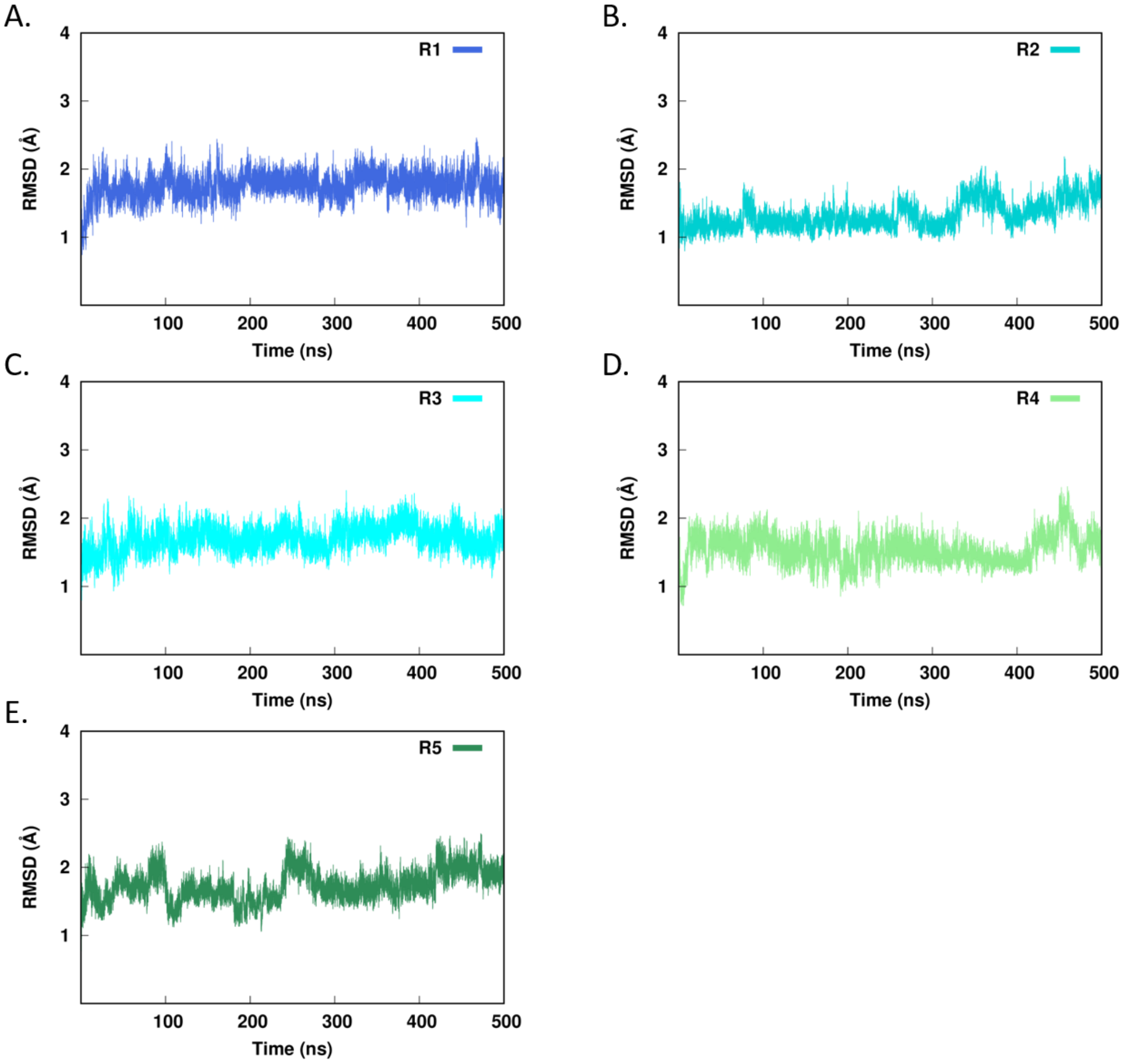


**Figure S6. Stability of Molecular Dynamics Simulations of Uncomplexed OXA-57.** Time series of backbone atom (N, Cα and C) RMSD values, compared to the starting structure, over 500 ns trajectories for repeat simulations one to five (R1-R5; panels A-E) of uncomplexed OXA-57.

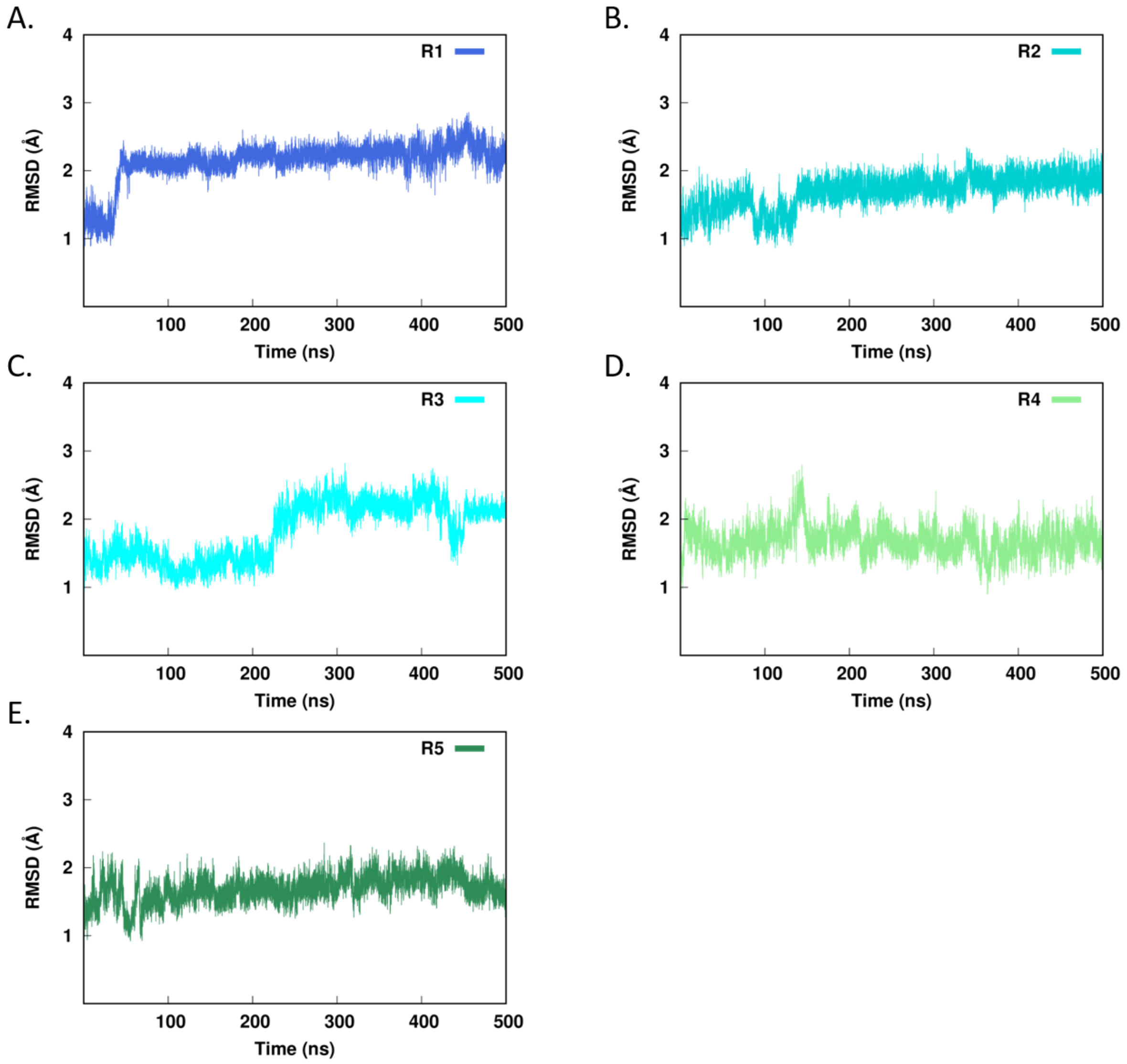


**Figure S7. Stability of Molecular Dynamics Simulations of OXA-57:Meropenem Complex** Time series of backbone atom (N, Cα and C) RMSD values, compared to the starting structure, over 500 ns trajectories for repeat simulations one to five (R1-R5; panels A - E) of the OXA-57:meropenem complex.

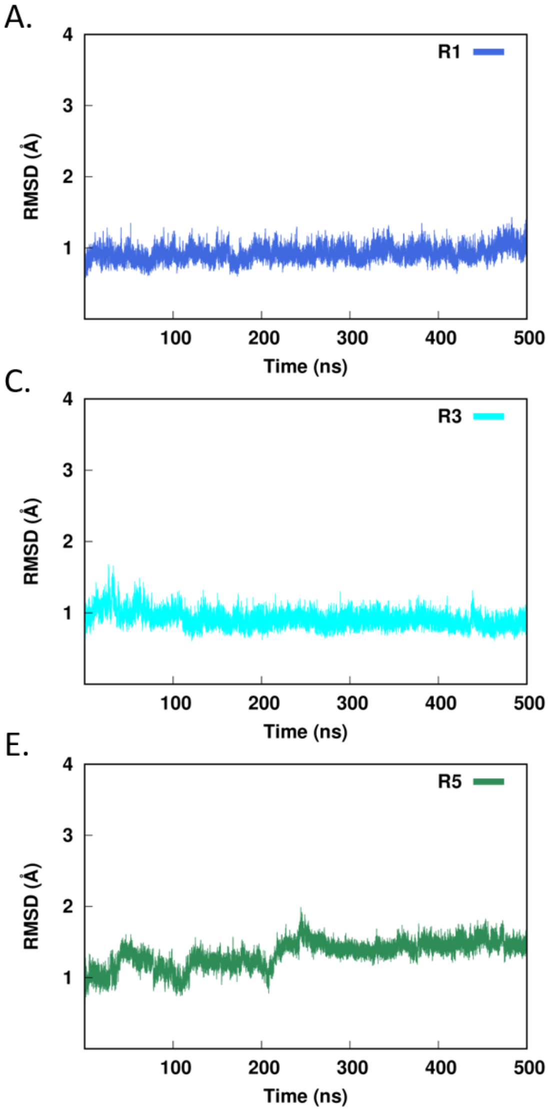


**Figure S8. Stability of Molecular Dynamics Simulations of Uncomplexed OXA-57 (Core Regions).** Time series of uncomplexed OXA-57 backbone atom (N, Cα and C) RMSD values, compared to the starting structure, over 500 ns trajectories for repeat simulations one to five (R1-R5; panels A-E), excluding residues corresponding to the β4 - β5 loop (S205 - S220) and the C-terminus (L266 - R269).

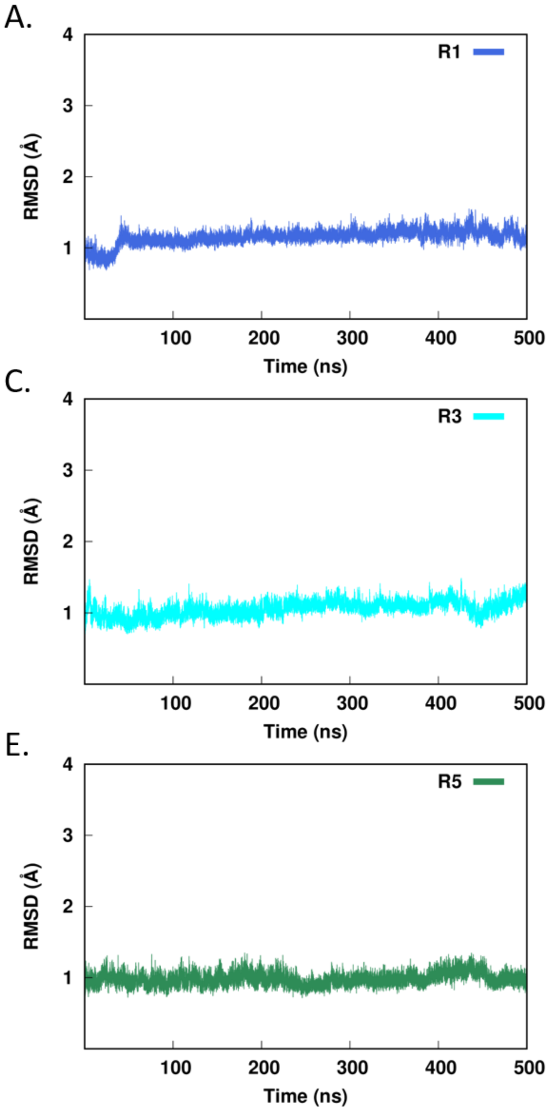


**Figure S9. Stability of Molecular Dynamics Simulations of OXA-57:Meropenem Complex (Core Regions).** Time series of backbone atom (N, Cα and C) RMSD values for the OXA-57:meropenem complex, compared to the starting structure, over 500 ns trajectories for repeat simulations one to five (R1 - R5; panels A - E), excluding residues corresponding to the β4 - β5 loop (S205 - S220) and the C-terminus (L266 - R269).

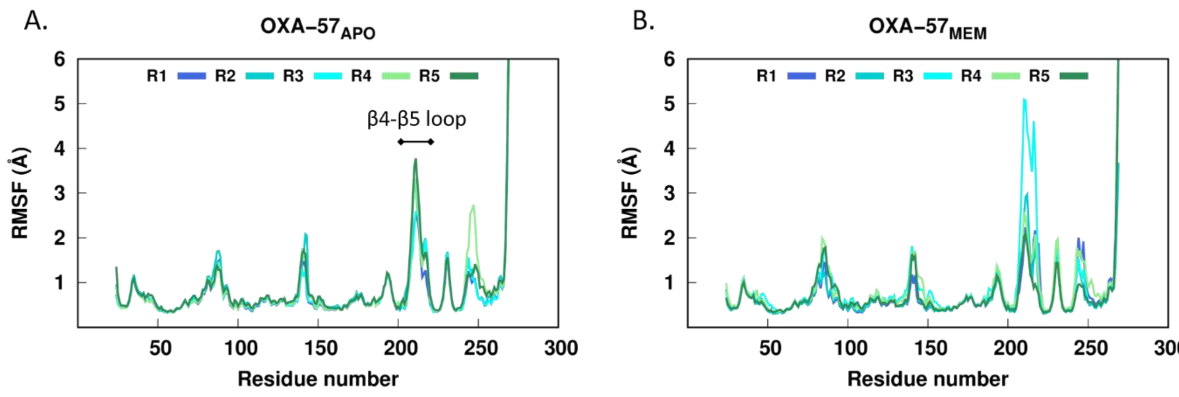


**Figure S10. Backbone Dynamics of OXA-57.** Backbone atom (N, Cα, C) RMSF values calculated over 425 ns MD trajectories (excluding first 75 ns for equilibration) using the average backbone (N, Cα, C) coordinates as reference for simulations of: A. uncomplexed OXA-57 and B. OXA-57:meropenem complex, respectively. The largest RMSF peak corresponds to the β4 - β5 loop.

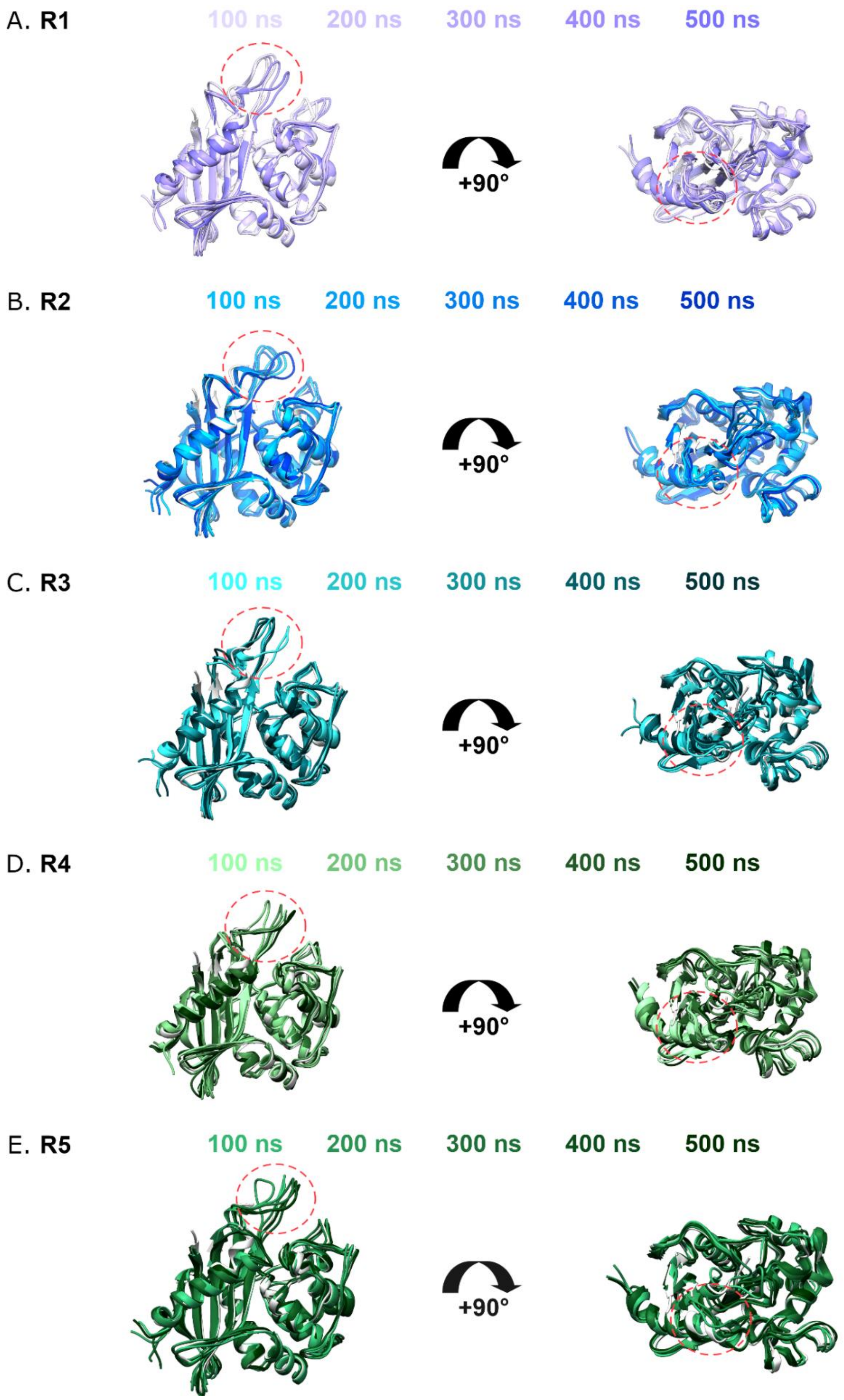


**Figure S11. Snapshots of Uncomplexed OXA-57 During MD Simulations.** Superposed structures from 100 ns snapshots throughout the 500 ns trajectories of simulations repeat one to five (R1-R5, A-E) of uncomplexed OXA-57. Coloring is as follows: R1 in purple, R2 in blue, R3 in cyan, R4 in mint and R5 in green. Data from each increasing 100 ns snapshot of the simulation is depicted as the repeat color becoming progressively darker. The crystal structure is shown in white and the β4-β5 loop is circled in red.

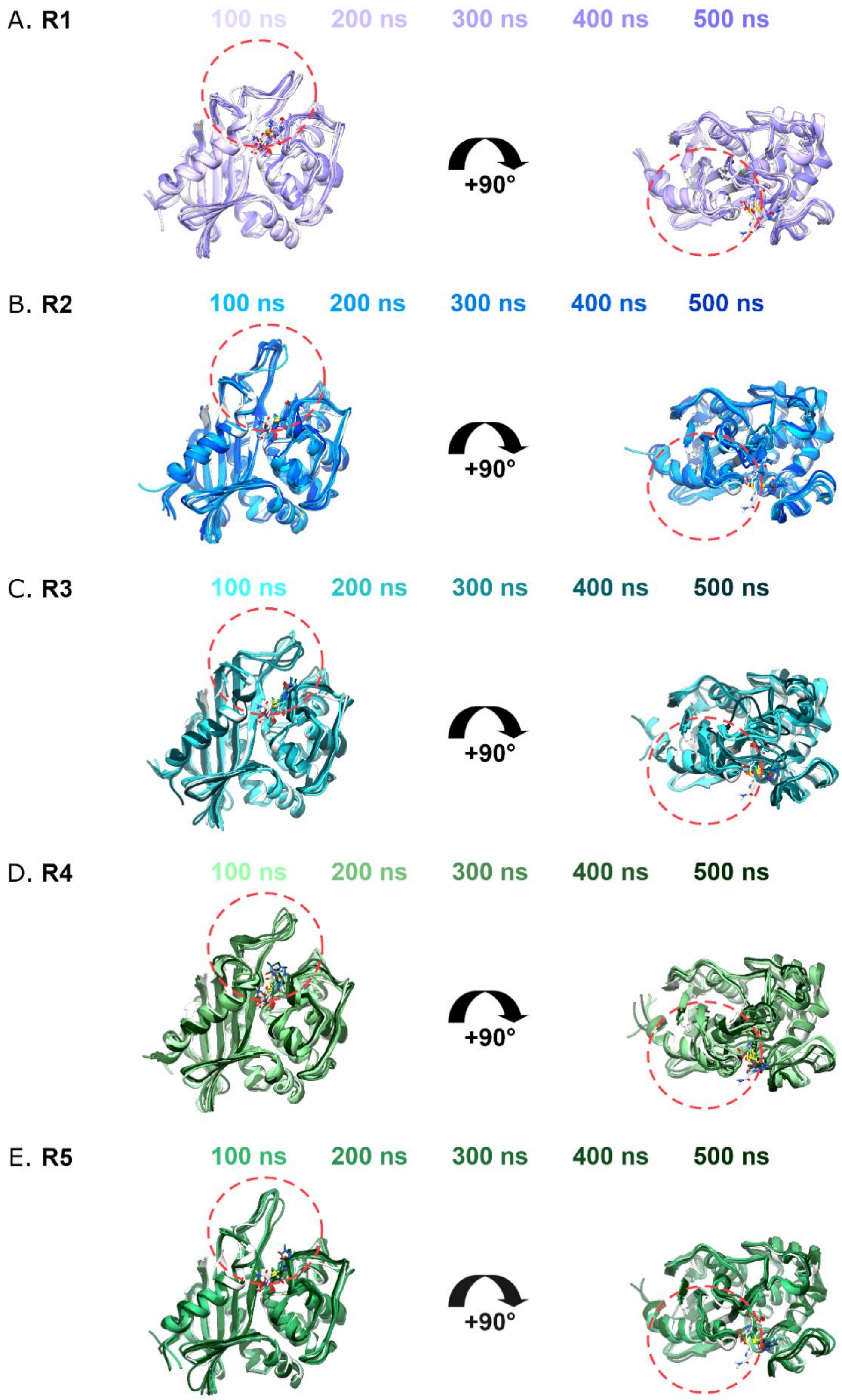


**Figure S12. Snapshots of OXA-57:Meropenem Complex During MD Simulations.** Superposed structures from 100 ns snapshots throughout the 500 ns MD trajectories of the OXA-57:meropenem complex for repeat simulations one to five (R1 - R5, panels A - E). Coloring is as follows: R1 in purple, R2 in blue, R3 in cyan, R4 in mint and R5 in green. Data from each increasing 100 ns snapshot of the simulation is depicted as the repeat color becoming progressively darker. The crystal structure is shown in white and the β4 - β5 loop is circled in red.

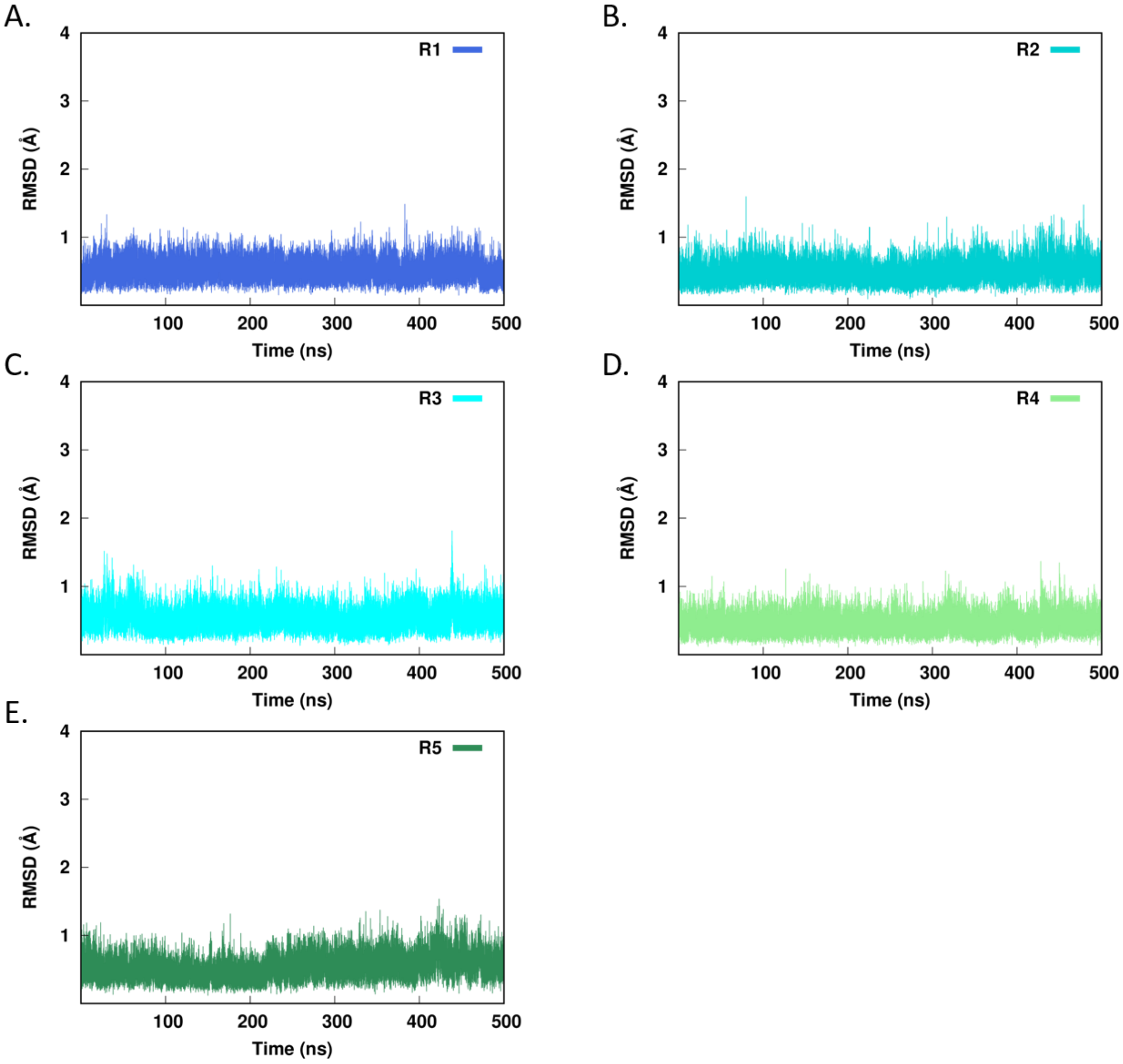


**Figure S13. Stability of Active Site Carbamyl-K56 During Molecular Dynamics Simulations of Uncomplexed OXA-57.** Time series of RMSD values excluding backbone atoms (N, Cα, C and O), compared to the starting structure, for the carbamylated active site lysine (K56), over 500 ns trajectories for repeat simulations one to five (R1 - R5; panels A - E) for uncomplexed OXA-57.

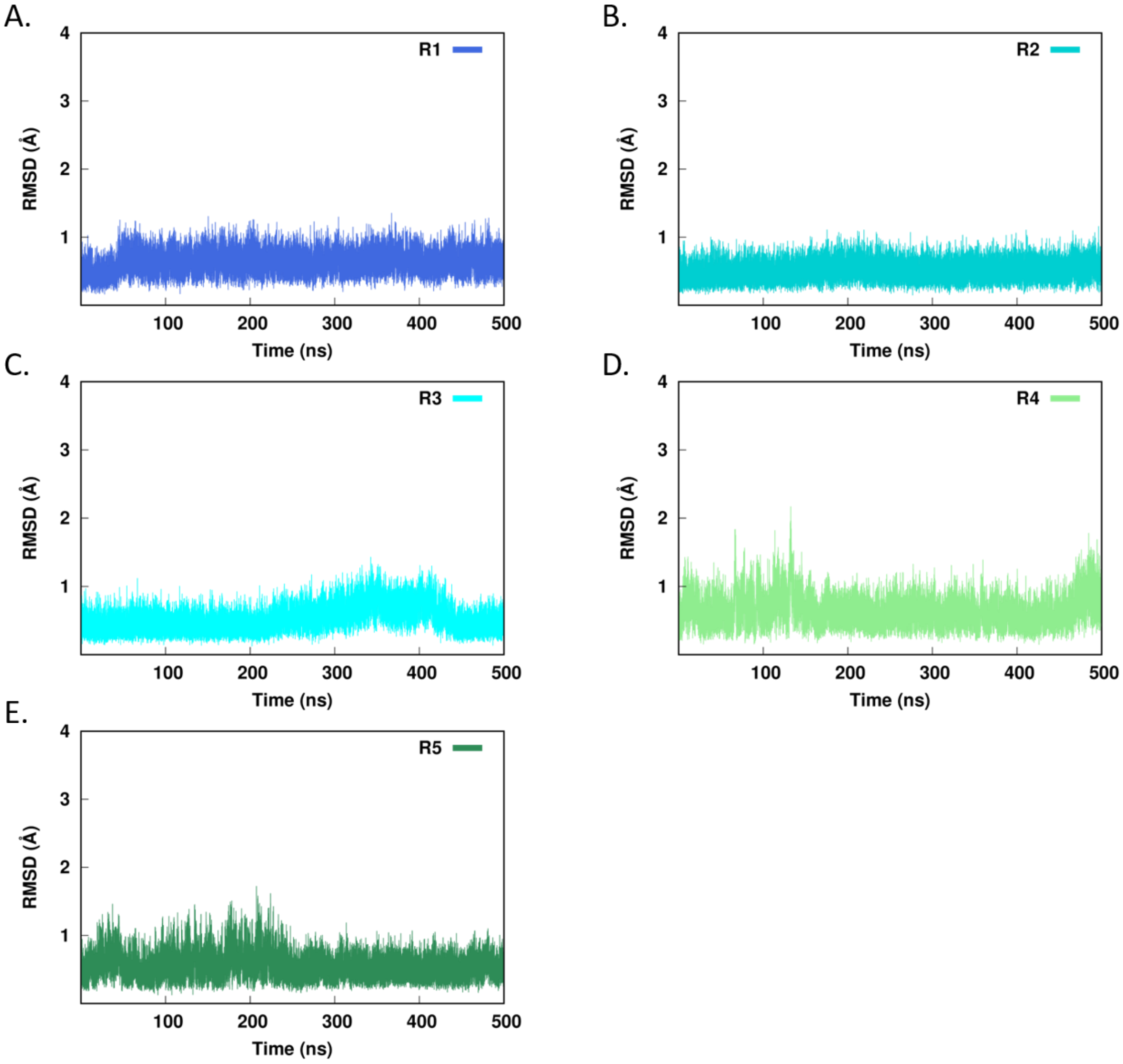


**Figure S14. Stability of Active Site Carbamyl-K56 During Molecular Dynamics Simulations of OXA-57:Meropenem Complex.** Time series of RMSD values excluding backbone atoms (N, Cα, C and O), compared to the starting structure, for the active site carbamylated lysine (K56) over 500 ns trajectories for repeat simulations one to five (R1 - R5; panels A - E) of the OXA-57:meropenem complex.

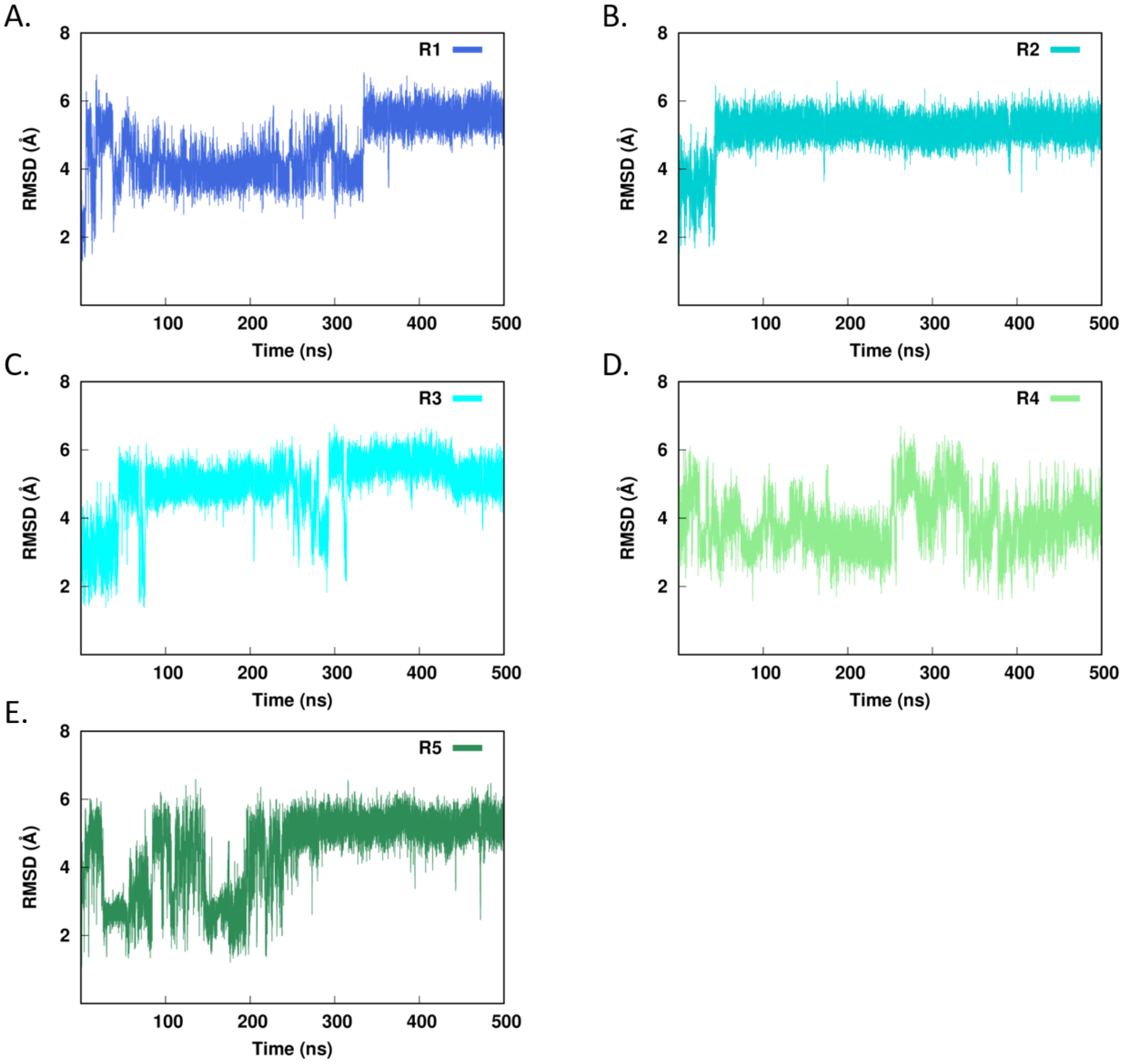


**Figure S15. Stability of Bound Meropenem During Molecular Dynamics Simulations of OXA-57:Meropenem Complex.** Time series of RMSD values, compared to the starting structure and excluding backbone atoms (N, Cα, C and O) of the acylated serine residue, for bound meropenem (MEM53) over 500 ns trajectories for repeat simulations one to five (R1 - R5; panels A-E) of the OXA-57:meropenem complex.

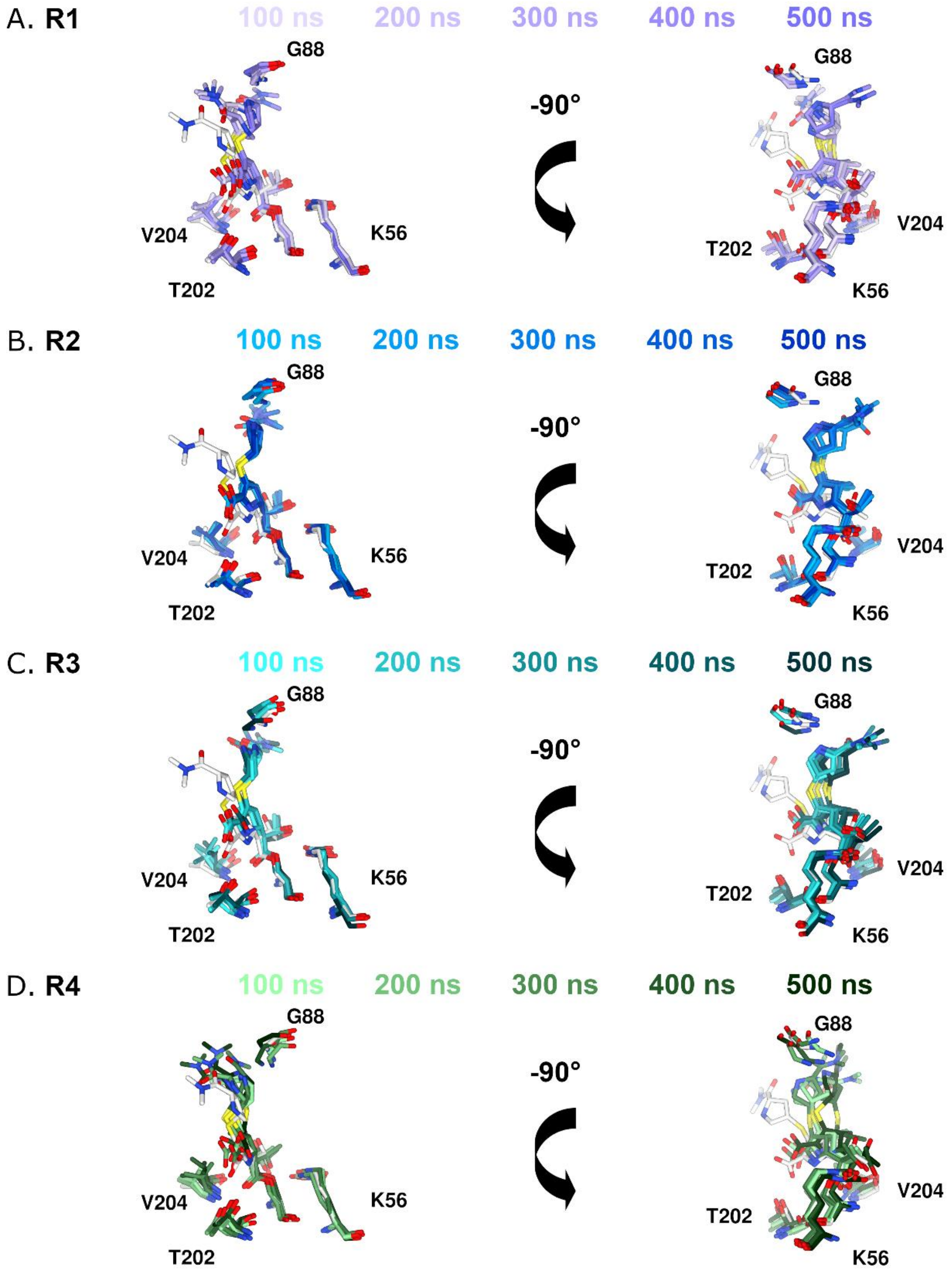


**Figure S16. Orientations of Bound Meropenem During MD Simulations.** Superposed structures of meropenem at 100 ns intervals throughout the 500 ns MD trajectories for repeat simulations one to four (R1-R4; panels A-D) of the OXA-57:meropenem complex. Coloring is as follows: R1 in purple, R2 in blue, R3 in cyan and R4 in mint. Data from each increasing 100 ns snapshot of the simulation is depicted as the repeat color becoming progressively darker. The crystal structure is shown in white.

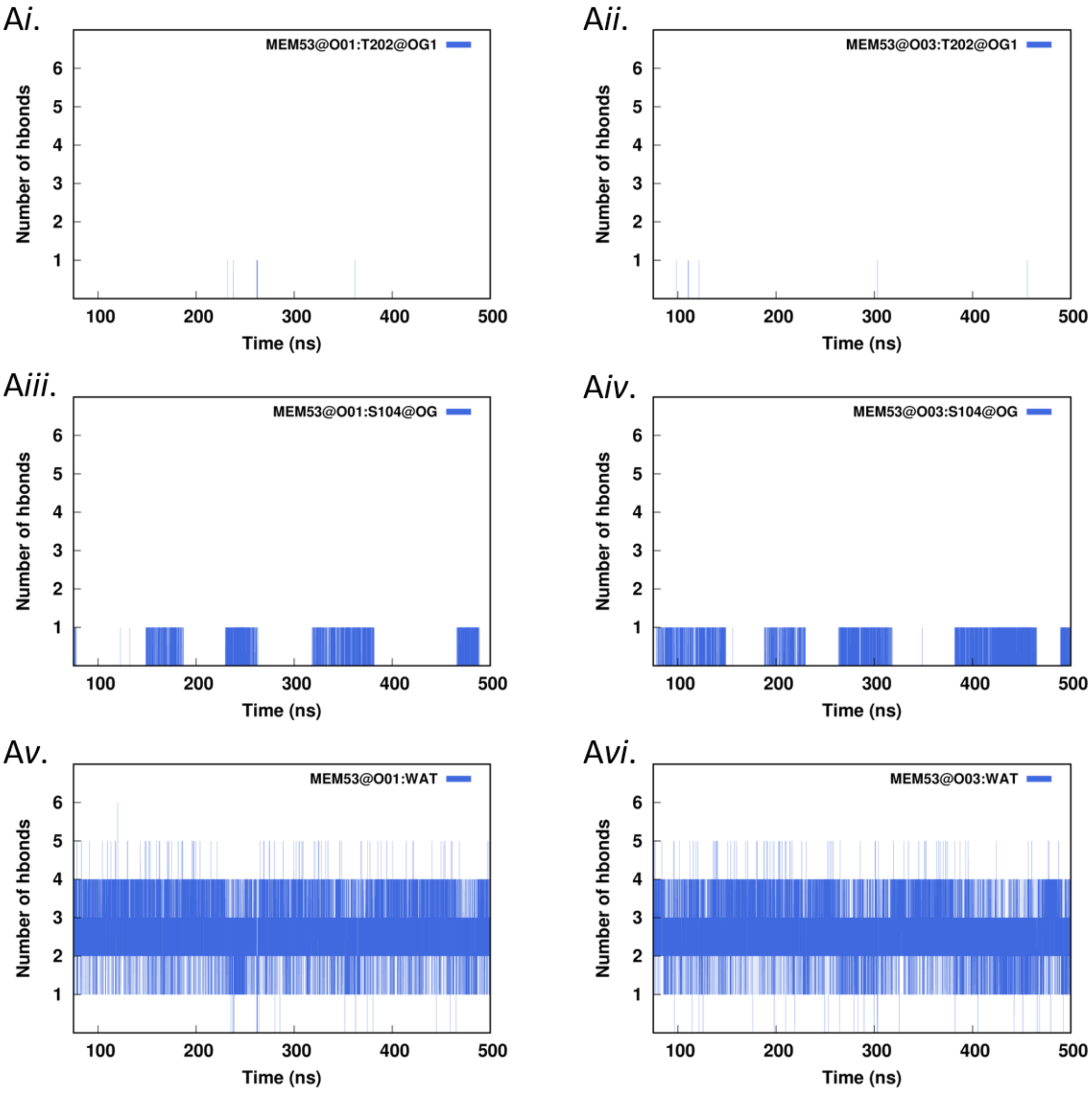
Ai.
MEM53@O01:T202@OG1
Aii.
MEM53@O03:T202@OG1
Aiii.
MEM53@O01:S104@OG
Aiv.
MEM53@O03:S104@OG
Av.
MEM53@O01:WAT
Avi.
MEM53@O03:WAT
Number of hbonds
1
2
3
4
5
6
Time (ns)
100
200
300
400
500

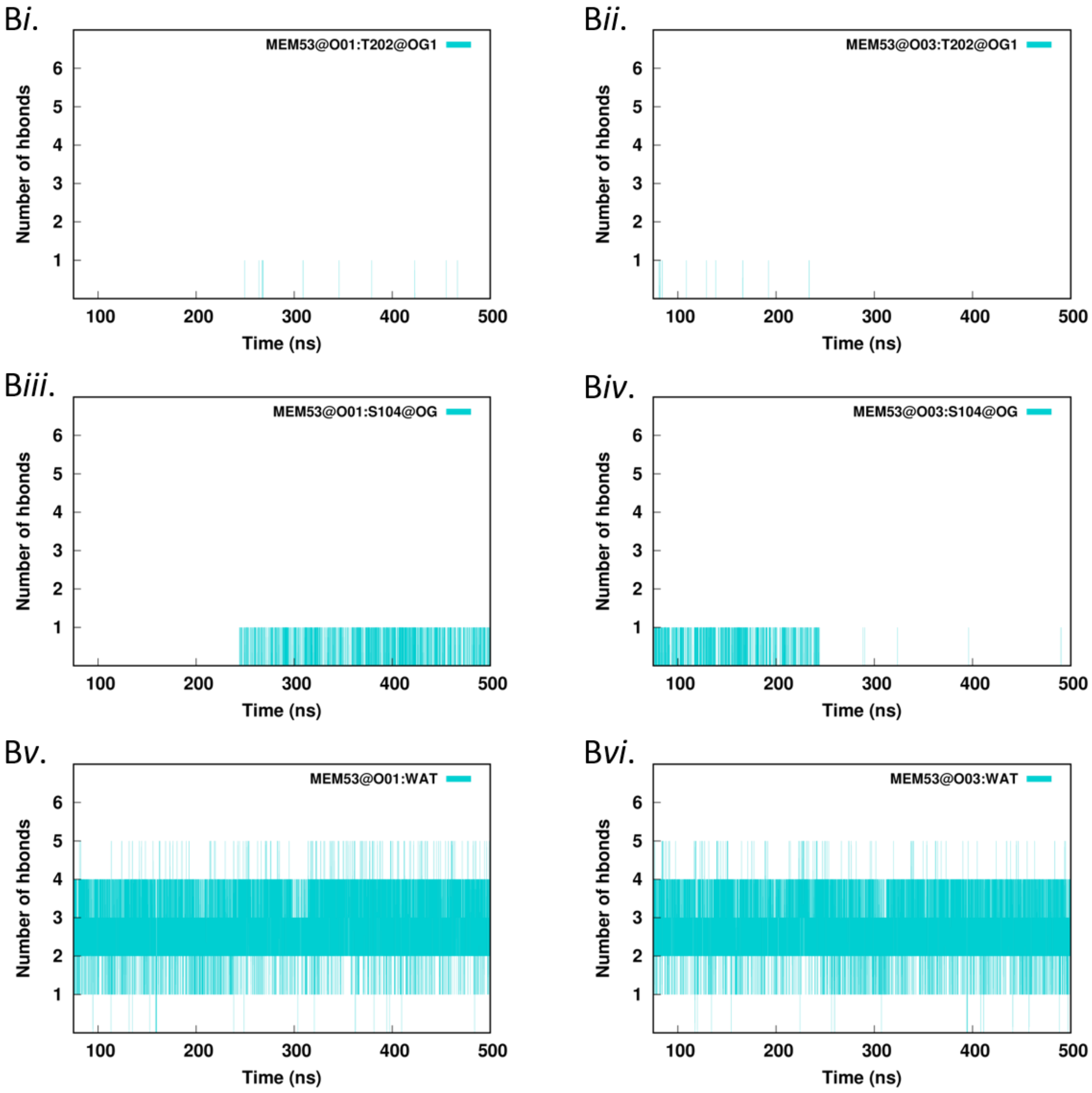

Bi.
MEM53@O01:T202@OG1
Bii.
MEM53@O03:T202@OG1
Biii.
MEM53@O01:S104@OG
Biv.
MEM53@O03:S104@OG
Bv.
MEM53@O01:WAT
Bvi.
MEM53@O03:WAT
Number of hbonds
Time (ns)
1
2
3
4
5
6
100
200
300
400
500

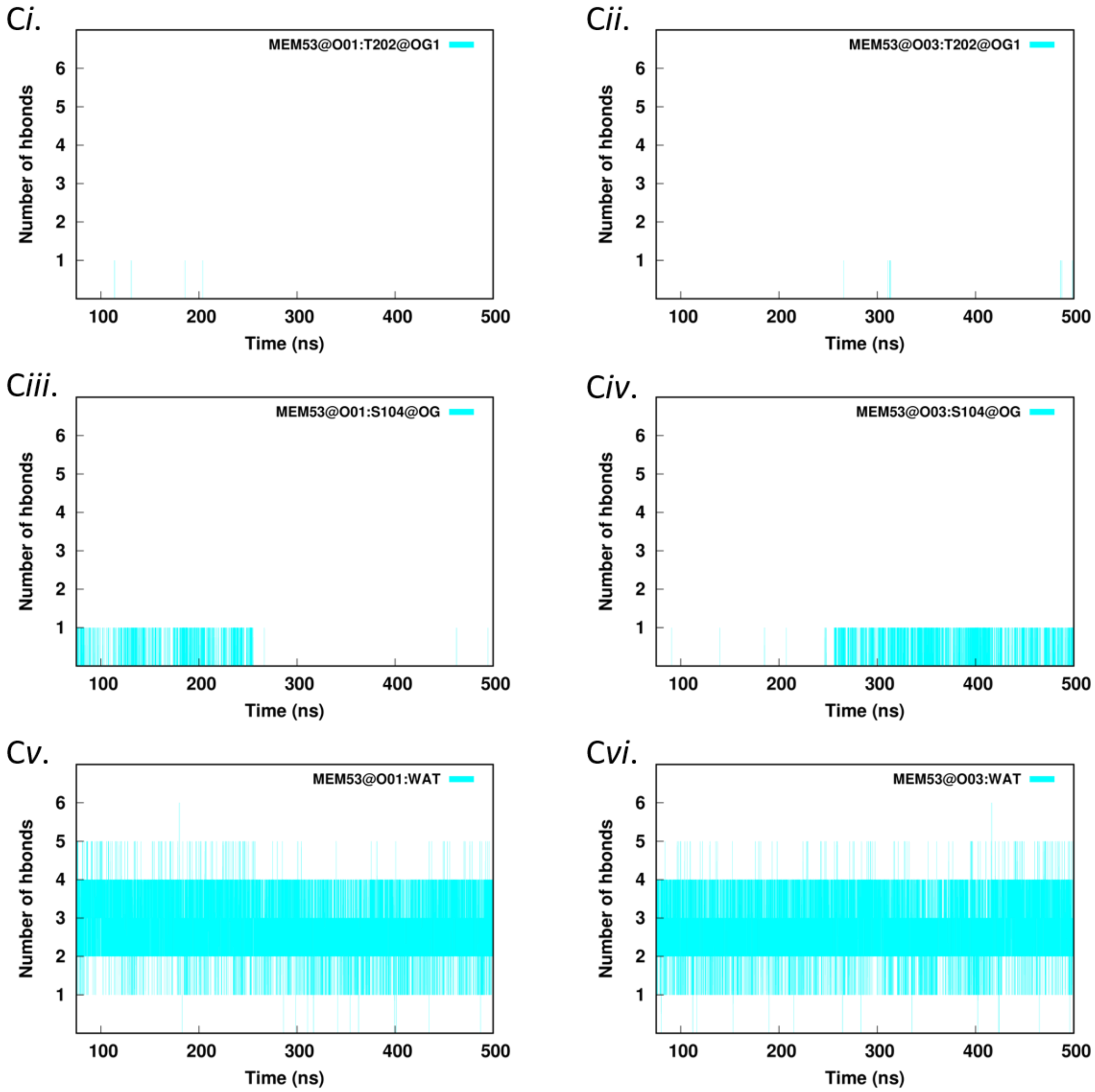
Ci.
MEM53@O01:T202@OG1
Cii.
MEM53@O03:T202@OG1
Ciii.
MEM53@O01:S104@OG
Civ.
MEM53@O03:S104@OG
Cv.
MEM53@O01:WAT
Cvi.
MEM53@O03:WAT
Number of hbonds
Time (ns)

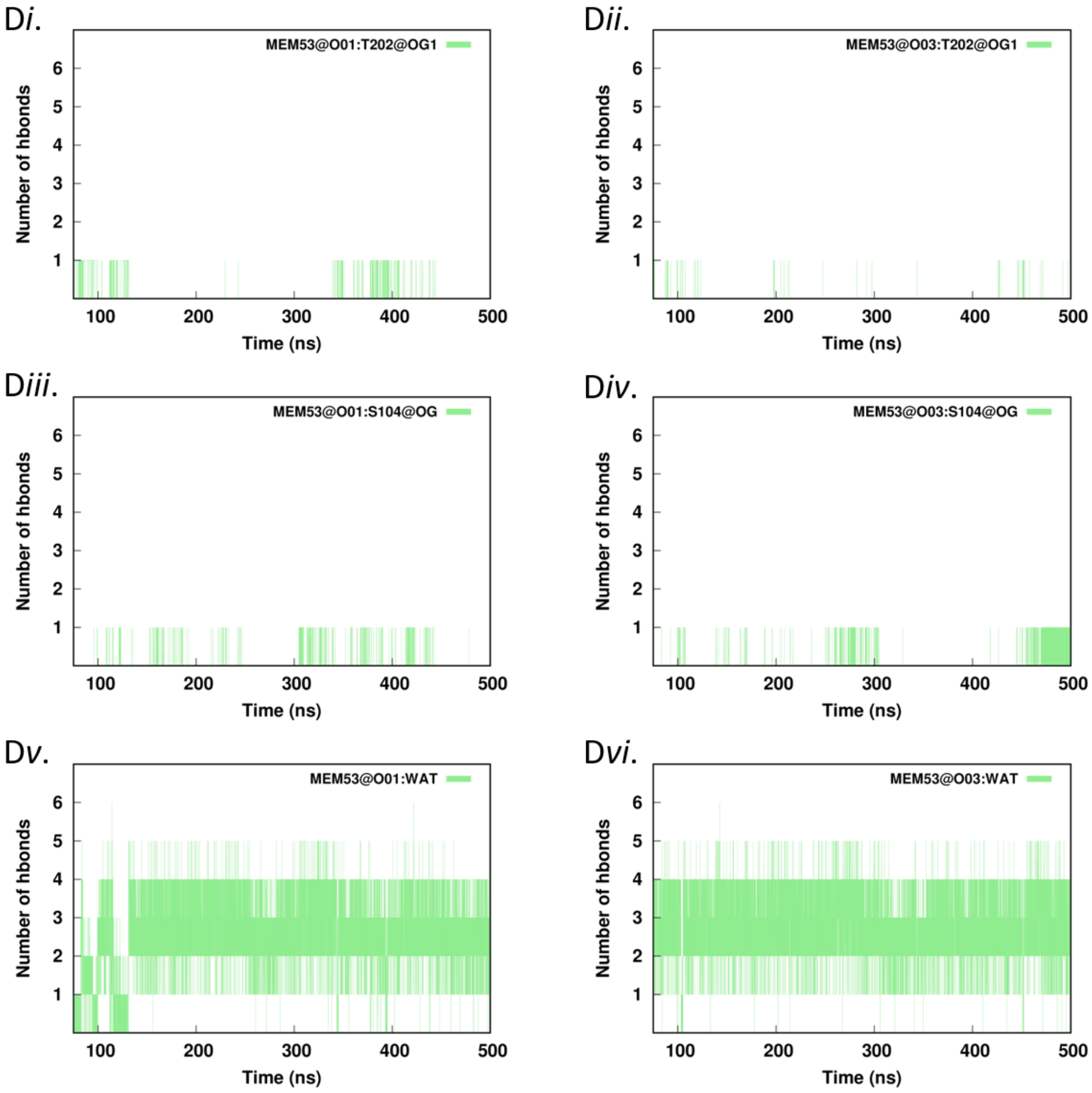

Di.
MEM53@O01:T202@OG1
Number of hbonds
Time (ns)
Dii.
MEM53@O03:T202@OG1
Number of hbonds
Time (ns)
Diii.
MEM53@O01:S104@OG
Number of hbonds
Time (ns)
Div.
MEM53@O03:S104@OG
Number of hbonds
Time (ns)
Dv.
MEM53@O01:WAT
Number of hbonds
Time (ns)
Dvi.
MEM53@O03:WAT
Number of hbonds
Time (ns)

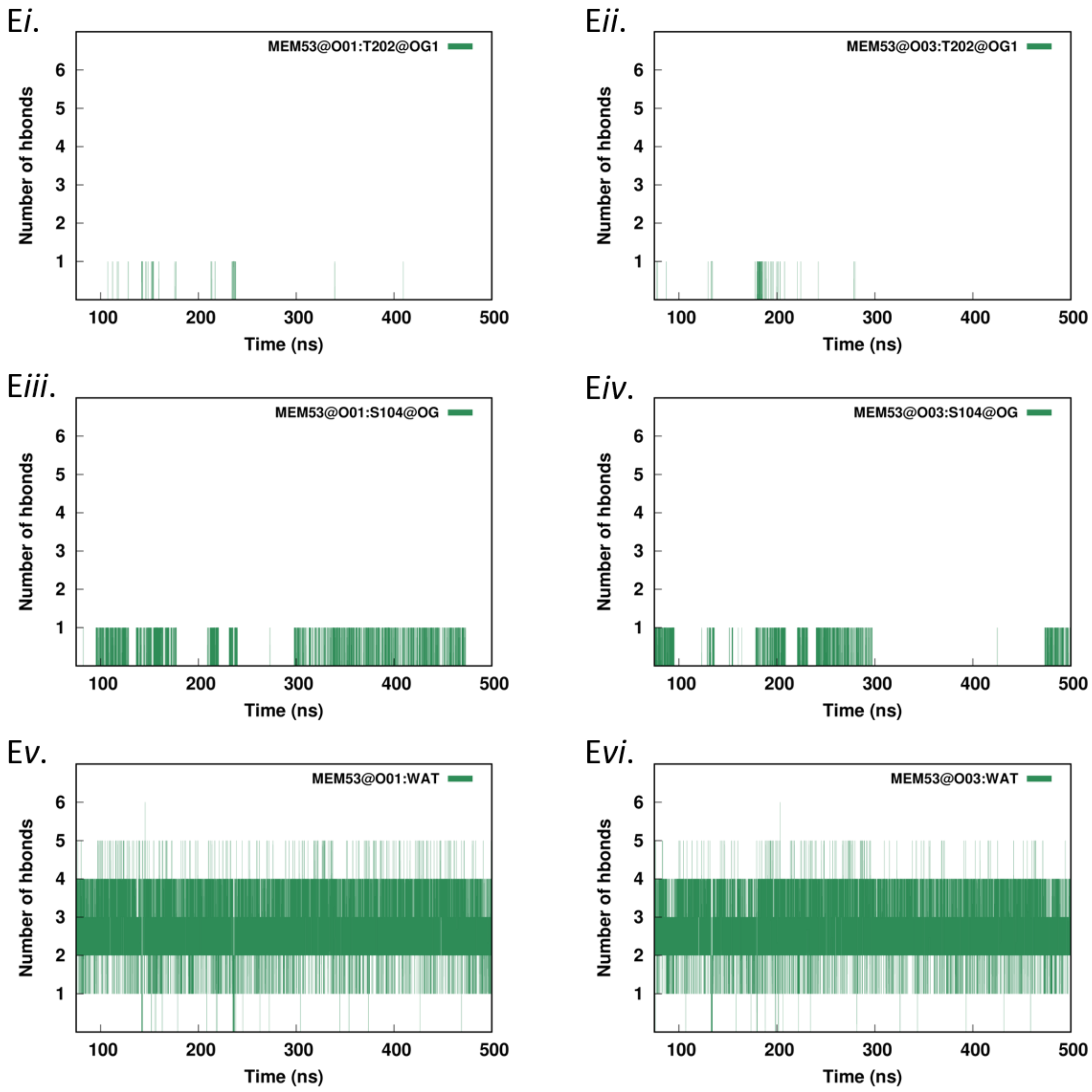


**Figure S17. Time-Dependence of Hydrogen Bonds during MD Simulations of the OXA-57:Meropenem Complex.** Time series of hydrogen bond interactions made by the meropenem carboxylate group through 425 ns MD trajectories of the OXA-57:meropenem complex simulations repeat one to five (R1-R5; panels A-E). (Residue number and atom name are given as acceptor:donor). *i*. MEM53@*O01*:T202@*OG1*, *ii*. MEM53@*O03*:T202@*OG1*, *iii*. MEM53@*O01*:S104@*OG*, *iv*. MEM53@*O03*:S104@*OG*, *v*. MEM53@*O01*:WAT, and *vi*. MEM53@*O03*:WAT. WAT = water molecules.

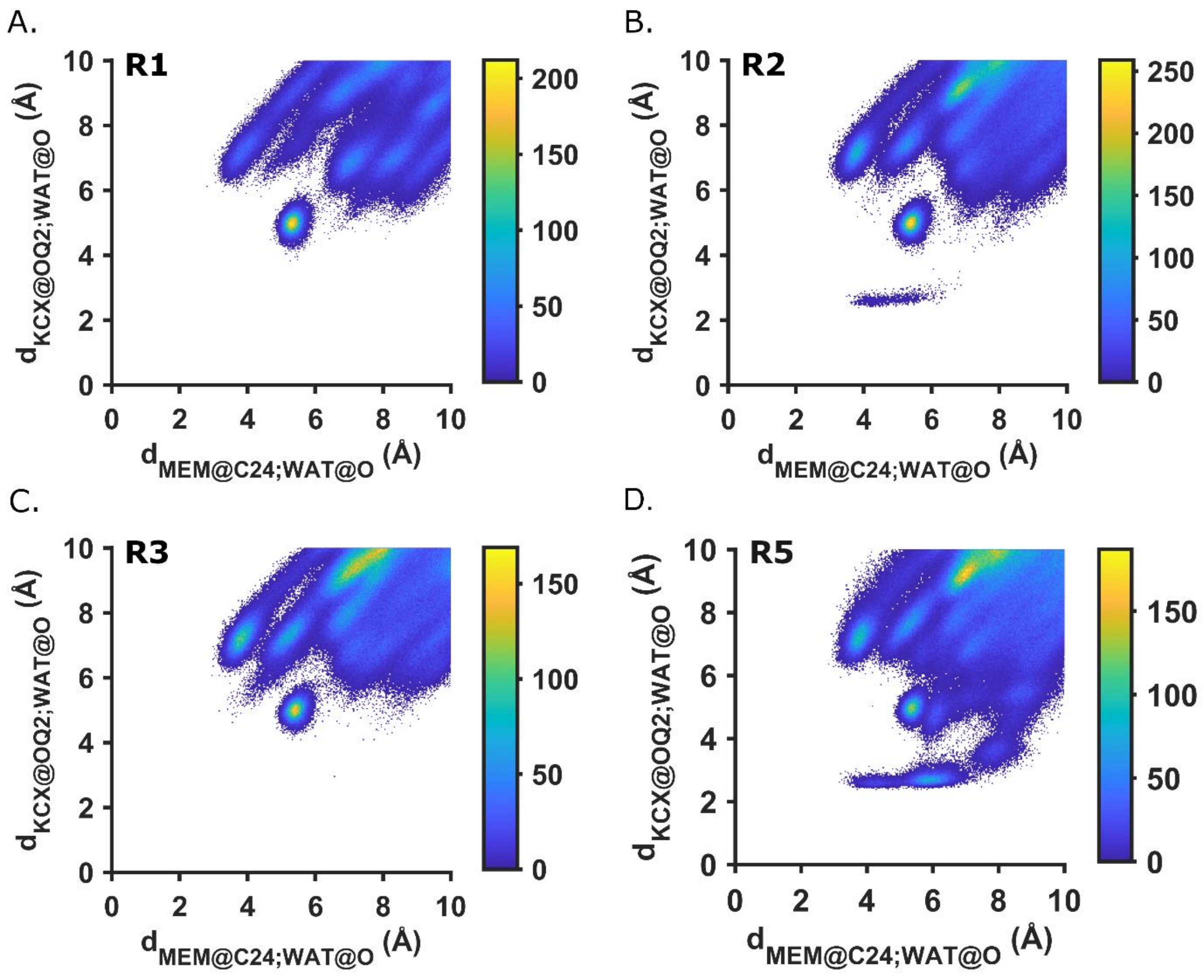


**Figure S18. Active Site Water Distribution during MD Simulations of the OXA-57:Meropenem Complex.** Distance distributions of water molecules between MEM53@*C24* and K56@*OQ2* over 425 ns MD trajectories of the OXA-57:meropenem complex in repeat simulations one, two, three and five (R1 - 3, R5; panels A - D).

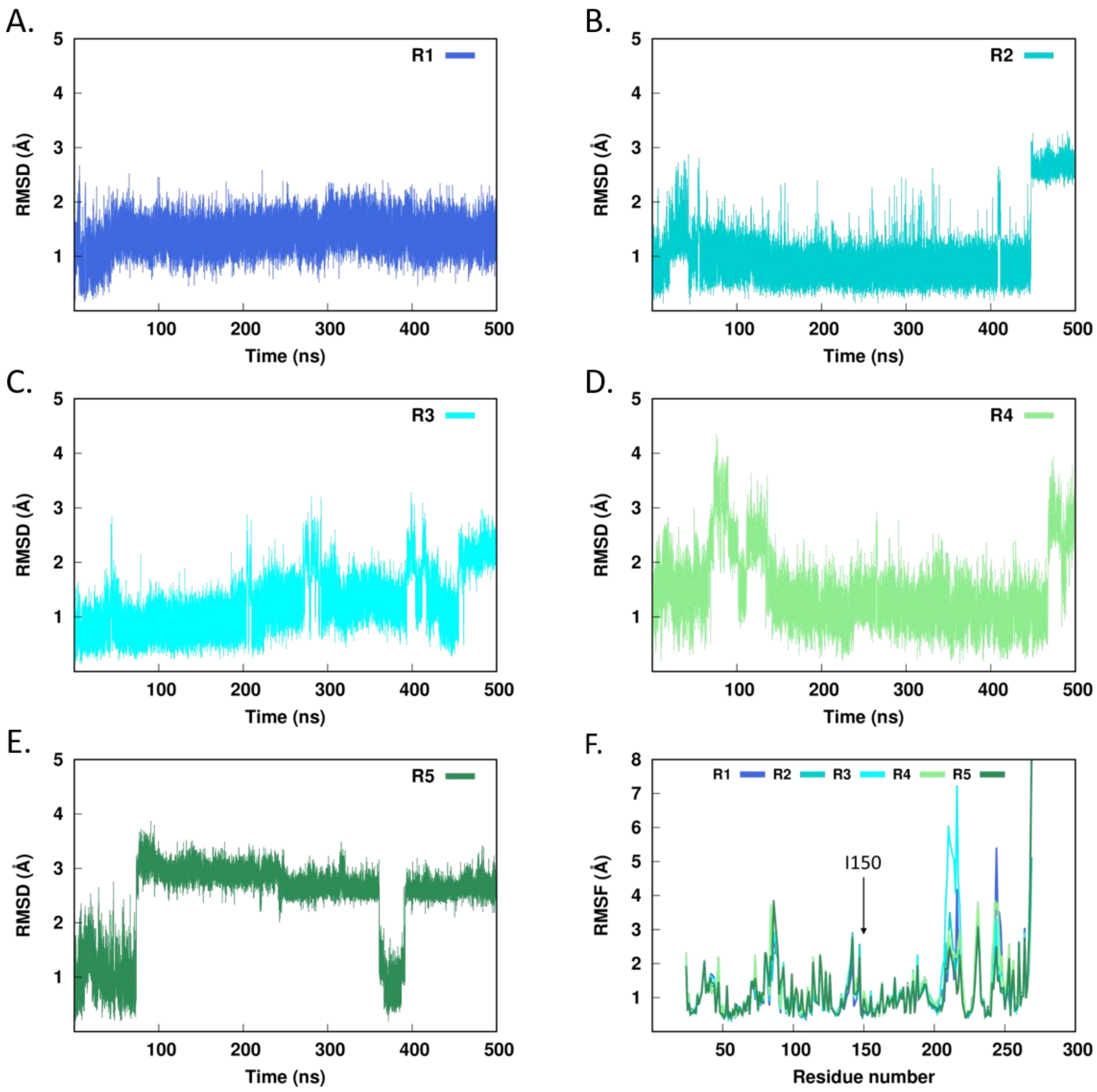


**Figure S19. Mobility of Residue I150 during MD Simulations of the OXA-57:Meropenem Complex.** RMSD distributions of the I150 sidechain (excluding atoms N, Cα, C, O, and hydrogens) for repeat simulations one to five (R1-R5; panels A-E) calculated over 425 ns trajectories of the OXA-57:meropenem complex. F. Per-residue RMSF distributions of all sidechains in the OXA-57:meropenem complex simulations. I150 is annotated with a black arrow.

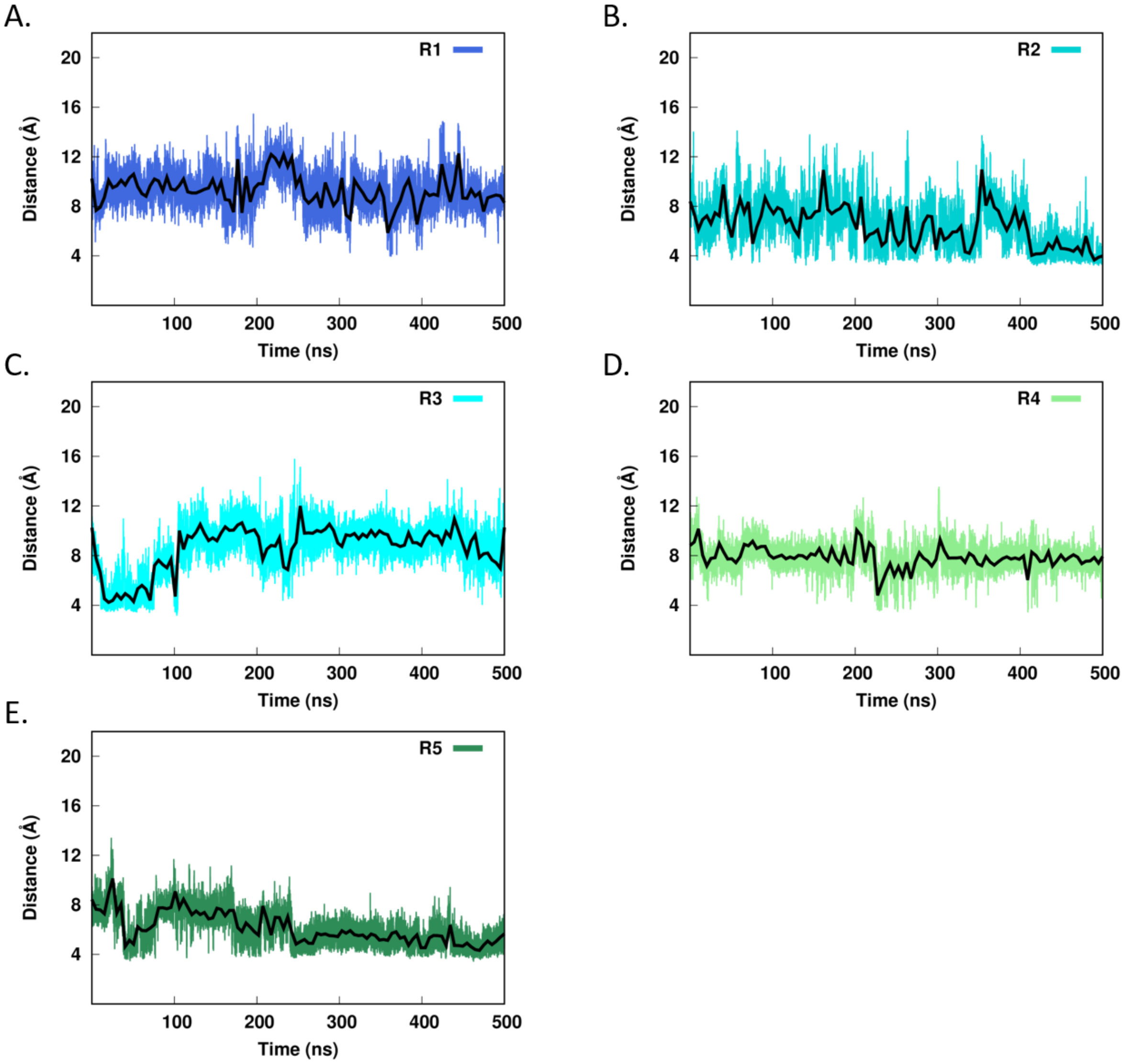


**Figure S20. Mobility of Active Site Loops during MD Simulations of Uncomplexed OXA-57.** Time series of the minimum distance between any carbon atom in residues W86 (α1 - α2 loop) and Y206 (β4 - β5 loop) calculated over 500 ns MD trajectories of uncomplexed OXA-57 for repeat simulations one to five (R1-R5; panels A-E). Black lines represent the distance data smoothed by the approximating cubic spline operation in gnuplot.

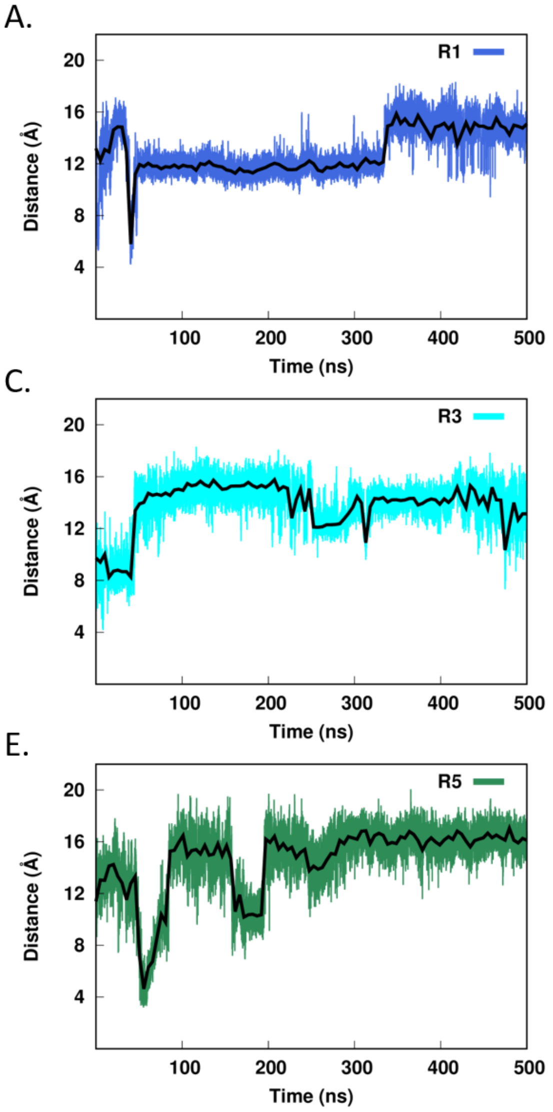


**Figure S21. Mobility of Active Site Loops during MD Simulations of the OXA-57:Meropenem Complex.** Time series of the minimum distance between any carbon atom in residues W86 (α1 - α2 loop) and Y206 (β4 - β5 loop) of the OXA-57:meropenem complex calculated over 500 ns MD trajectories for repeat simulations one to five (R1 – R5; panels A-E). Black lines represent the distance data smoothed by the approximating cubic spline operation in gnuplot.

A.

B.

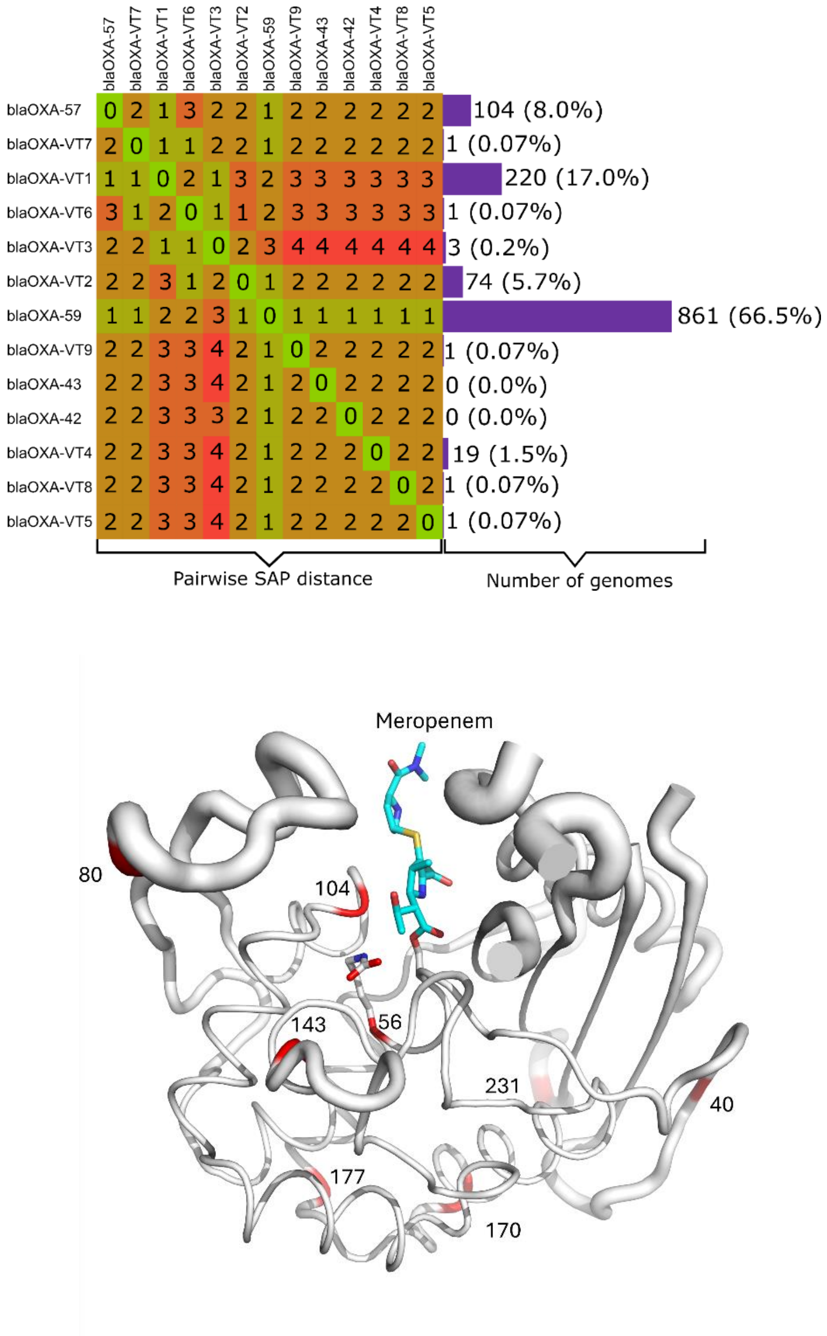


**Figure S22. Sequence Polymorphisms in *B. pseudomallei* Chromosomal OXA β-Lactamases**. A. Pairwise single amino acid polymorphism (SAP) distances (numbers of residue substitutions) of previously known blaOXA isoforms (OXA-42, OXA-43, OXA-57, OXA-59)) and new variant types (VTs, 1-9) identified in genomes of 1,295 clinical *B. pseudomallei* isolates collected in Northeast Thailand. Numbers (right) denote absolute numbers and percentage frequency of occurrence of individual isoforms. Sequences are deposited in the European Nucleotide Archive (ENA) under study accession numbers PRJEB25606 and PRJEB35787. B. Positions of residue substitutions present in known and newly identified OXA isoforms (red) mapped onto crystal structure of OXA-57:meropenem complex (PDB 9HPW, this work). Bound meropenem (carbon atoms cyan) and the side chains of residues S53 and K56 (carbon atoms white) are shown as sticks. Worm thickness indicates B-factor. This image was created in Pymol (www.pymol.org).

(22) Langley, G. W.; Cain, R.; Tyrrell, J. M.; Hinchliffe, P.; Calvopiña, K.; Tooke, C. L.; Widlake, E.; Dowson, C. G.; Spencer, J.; Walsh, T. R.; Schofield, C. J.; Brem, J. Profiling Interactions of Vaborbactam with Metallo-β-Lactamases. *Bioorg. Med. Chem.*

*Lett.* **2019**, *29* (15), 1981–1984. https://doi.org/10.1016/J.BMCL.2019.05.031.

(23) Tsivkovski, R.; Totrov, M.; Lomovskaya, O. Biochemical Characterization of QPX7728, a New Ultrabroad-Spectrum Beta-Lactamase Inhibitor of Serine and Metallo-Beta-Lactamases. **2020**. https://doi.org/10.1128/AAC.
